# FoxO1 and HNF1α cooperatively remodel nucleosome architecture and mechanical stability to activate transcription

**DOI:** 10.64898/2026.11.01.763477

**Authors:** Alfredo Jose Florez Ariza, Zhenlin Yang, Robert P. Sosa, César Díaz-Cellis, Hannah J. Richter, Vijay Ramani, Carlos Bustamante, Eva Nogales

## Abstract

How different transcription factors (TFs) cooperate to bind nucleosomal DNA targets remains unclear. Cryo-EM and force spectroscopy show that the pioneer factor FoxO1 binds superhelical location (SHL) +2 of nucleosomes assembled on the human IGFBP1 promoter, enhances distal DNA unwrapping, destabilizes the histone core, and reduces nucleosome mechanical stability. TF HNF1α binds SHL +5/+7 as a dimer, induces extensive DNA unwrapping without perturbing the histone core, and reshapes nucleosome disassembly dynamics. FoxO1 enhances HNF1α recruitment, and together they stabilize the nucleosome during its mechanically-induced disassembly trajectories. In-cell nucleosome footprinting on active promoters shows that FoxO1 binding enhances unwrapped nucleosomes and subnucleosomes presence, while HNF1α increases stabilized and “breathing” nucleosomes and reduces the subnucleosome population. Combined, they display an intermediate nucleosome phenotype. Our data leads to a model where FoxO1 destabilizes the nucleosome and enhances HNF1α recruitment, and together they remodel nucleosome architecture both in vitro and in native chromatin.

## INTRODUCTION

In eukaryotes, genomic DNA is packaged into chromatin, whose fundamental unit is the nucleosome. During processes such as transcription or DNA replication, chromatin-associated factors influence chromatin structure, dynamics, and higher order organization^4–6^. Structural studies depict the nucleosome as a stable assembly of 147 bp of DNA wrapped around a histone octamer that restricts access of transcription factors (TFs) to most of the nucleosomal DNA^1–3^. Studies using single molecule optical tweezers have used controlled mechanical perturbation on nucleosomes, which can be considered to mimic the type of action exerted by chromatin remodelers in vivo, to probe nucleosomes mechanical properties, showing that nucleosomes act as barriers to transcription, and that their mechanical stability can be modulated by histone variants, post-translational modifications, and transcriptional regulators^4,5^. Understanding how TFs overcome the nucleosomal mechanical barrier is central to deciphering how gene expression is regulated in a chromatin context. Pioneer factors constitute a specialized class of TFs that can recognize their target sequences within nucleosomal DNA in heterochromatin regions^6,7^, initiating nucleosome rearrangements that enable subsequent access by additional factors, thereby playing an important role in the regulation of gene expression during cell reprogramming and differentiation^6,8–11^.

FoxO1 is a member of the Forkhead box O (FoxO) family of pioneer factors^12^ that is highly expressed in insulin-responsive tissues, where it regulates genes involved in metabolism and apoptosis, and its dysregulation has been associated with cancer progression^12–15^. In humans, a FoxO1 binding site is located within the proximal promoter of the insulin-like growth factor–binding protein 1 (IGFBP1) gene, a member of the carrier proteins family highly expressed in hepatocytes. IGFBP1 modulates the bioavailability of insulin-like growth factor proteins (IGFs) involved in regulation of cell growth, proliferation and differentiation^16–19^. Expression of IGFBP1 is also regulated by the hepatocyte nuclear factor 1α (HNF1α), a homeobox canonical TF essential for pancreatic β-cells function and hepatocyte differentiation^20,21^. Importantly, mutations of this TF are associated with early-onset diabetes^22–24^. Of note, *in vitro* and *in vivo* studies have shown that both FoxO1 and HNF1α are directly involved in chromatin opening^14,25–27^. Crystal structures have shown how each of these factors bind to linear DNA targets^22,28–30^, but how these binding modes are accommodated within a chromatin context or how they affect each other’s function is unknown. Thus, the molecular mechanisms by which FoxO1 and HNF1α could jointly elicit structural rearrangements in nucleosomes that may affect their stability and the accessibility of the nucleosomal DNA remain elusive.

We have combined single-particle cryo-electron microscopy (cryo-EM) and dual-trap high-resolution optical tweezers force spectroscopy to elucidate the structural and mechanical basis of nucleosome remodeling by FoxO1 and HNF1α in vitro. Our studies reveal that FoxO1 destabilizes the nucleosome core to promote DNA unwrapping and enhances HNF1α recruitment, and together these factors cooperate to remodel nucleosome architecture and mechanics. In vivo nucleosome footprint assays indicate that FoxO1 favors an enhanced presence of unwrapped nucleosomes and subnucleosome species, while HNF1α is associated with the highest presence of stable and ‘breathing’ nucleosomes with accessible intra-nucleosomal DNA. Simultaneous binding of both factors leads to an intermediate scenario, where stable wrapped nucleosomes are enriched in comparison to the FoxO1 alone, but are less populated than when HNF1α alone is present. Together, our results show that FoxO1 and HNF1α cooperate to remodel the architecture and stability of IGFBP1 nucleosomes in vitro and in native chromatin in promoter regions.

## RESULTS

### The IGFBP1 nucleosome undergoes asymmetric ‘breathing’

To elucidate the molecular mechanisms by which FoxO1 and HNF1α bind to and affect nucleosome architecture, we reconstituted nucleosomes using recombinant *Xenopus laevis* histone octamers and a 158 bp DNA fragment corresponding to the human IGFBP1 proximal promoter (positions −215 to −58). This sequence contains an insulin-responsive element (IRE), which serves as the FoxO1 binding site, as well as a cognate binding site for HNF1α (Figure 1a and 1b and Figure S1A). Using cryo-EM, we revealed that IGFBP1 nucleosomes (^IGFBP^Nuc) exist in two major conformations, referred to as Class I (3.4 Å overall resolution) and Class II (3.7 Å overall resolution), with approximately equal abundance (Figure S3, Figure S8 and Table S1). Class I shows fully wrapped nucleosomal DNA and closely resembles the canonical, well-positioned Widom 601 nucleosome^1,2^. We unambiguously assigned the DNA register and orientation within the density map, localizing the FoxO1 binding site (IRE motif) near SHL +2 and the HNF1α binding site between SHLs +5 and +7 (Figure 1c). In contrast, Class II displays asymmetric DNA unwrapping. While the proximal DNA arm at the 5′ end of the IGFBP1 sequence (SHL −7) remains fully wrapped like in Class I, the distal DNA arm on the 3′ end exhibits approximately 8 bp of unwrapped DNA, with the last histone–DNA contact occurring near SHL +6 (Figure 1d). This local DNA unwrapping is accompanied by the loss of histone–DNA contacts involving the H3 N-terminal region, which is not resolved in the density map, consistent with increased conformational flexibility upon DNA release (Figure 1d). Our cryo-EM observations are consistent with the higher GC content and lower predicted curvature of the distal DNA arm (Figure S1b), which would exhibit a higher propensity to spontaneous unwrapping as shown by previous atomic force microscopy studies^31^.

**Figure 1:**
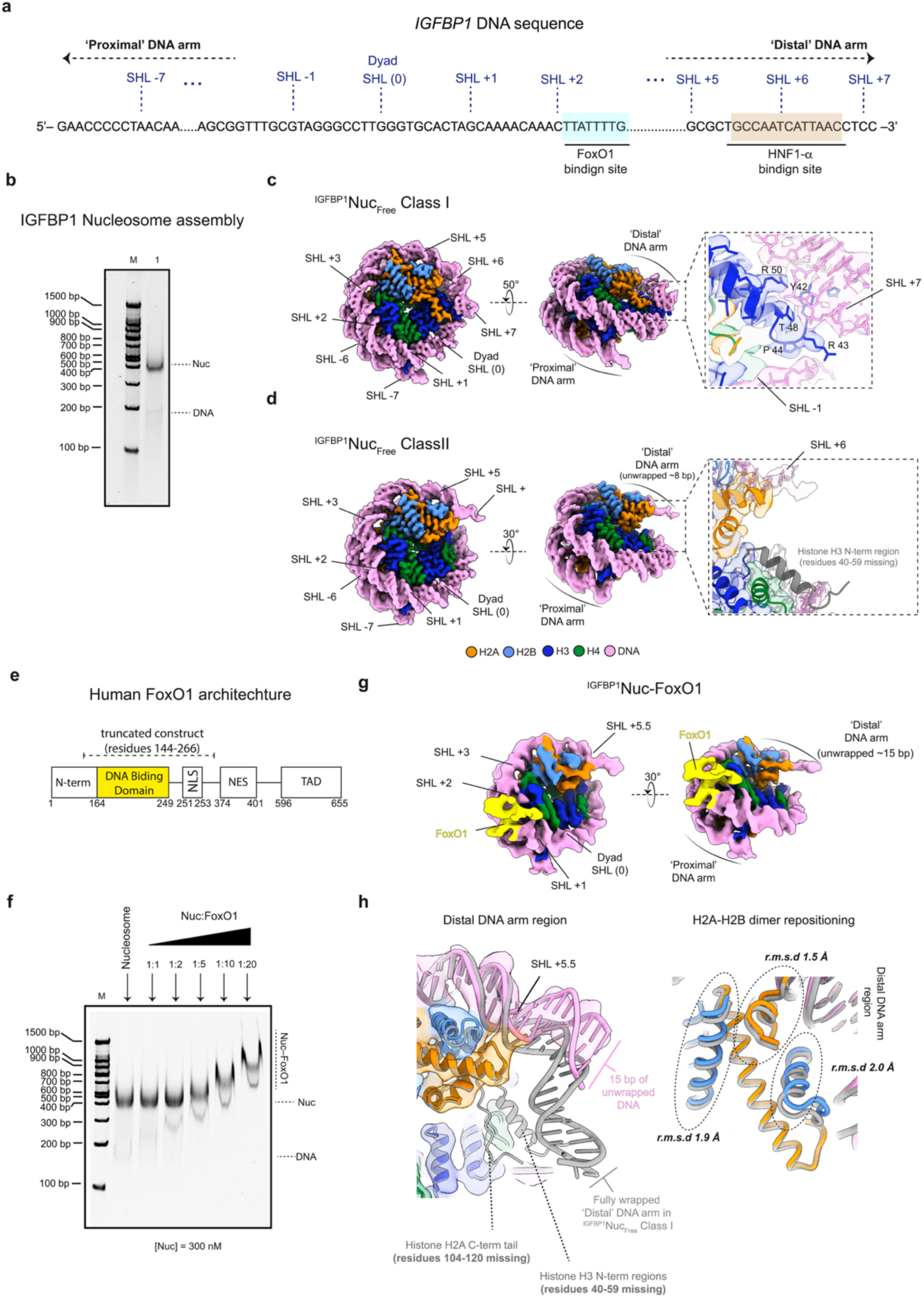
Cryo-EM structure of the IGFBP1 Nucleosome, alone (^IGFBP1^Nuc Class I and Class II) or bound to FoxO1 (^IGFBP1^Nuc–FoxO1). **a.** Schematic representation of the 158 bp *IGFBP1* sequence (-215 to -58). The binding sites for the FoxO1 and HNF1α are indicated. Superhelical locations (SHLs) relative to the nucleosome structure are indicated. The ‘proximal’ DNA arm is located on the 5’-end of the sequence, while the ‘distal’ DNA arm is located on the 3’-end. **b.** Native Tris base, boric acid, and EDTA (TBE) gel TBE (5%) showing the optimized assembly of the IGFBP1 nucleosome using an octamer:DNA molar ration of 1:0.9. **c.** Cryo-EM map of the fully wrapped, IGFBP1Nuc Class I structure shown in different views. The close-up view shows contacts between residues in the histone H3 N-terminal region and the distal DNA. Color key used is shown. **d.** Cryo-EM map of the partially unwrapped, *^IGFBP1^*Nuc Class II structure shown in different views. The distal DNA arm is unwrapped by ∼8 bp. The close-up view shows that residues in the H3 N-terminal region visible in **b** are not observed upon DNA unwrapping. **e.** Schematic representation of the human FoxO1 domain architecture. DNA binding domain (DBD), Nuclear Localization Sequence (NLS), Nuclear Export Sequence (NES), and Transactivation Domain (TAD). The truncated construct used in this work (residues 144-266) is indicated. **f.** Representative EMSA (5% TBE gel) for different nucleosome:FoxO1 molar ratios. Nucleosome concentration ([Nuc]) was kept constant at 300 nM. Bands corresponding to free DNA (DNA), nucleosome (Nuc), and Nucleosome bound to FoxO1 (Nuc–FoxO1), are indicated. **g.** Cryo-EM structure of the ^IGFBP1^NucFoxO1 complex shown in two orientations, with FoxO1 bound at ∼SHL +2. The distal DNA appears unwrapped by ∼15 bp. **h.** Left: Close-up view of the distal DNA region in the ^IGFBP1^Nuc_FoxO1_ map with fitted atomic model. The model of the fully wrapped *^IGFBP1^*Nuc Class I structure (gray ribbon) has been superimposed for reference. In the ^IGFBP1^Nuc_FoxO1_ structure, the last histone-DNA contact is mapped at the SHL 5.5, with ∼15 bp of the unwrapped distal DNA visible. The histone H2A C-term tail and H3 N-terminal region are not observed. Right: Close-up view of the distal H2A-H2B histone dimer in the ^IGFBP1^Nuc_FoxO1_ structure (color coded as indicated), with the *^IGFBP1^*Nuc Class I (gray ribbon) superimposed to show repositioning (r.m.s.d range between ∼1.5Å and ∼2Å)

### FoxO1 binds the IGFBP1 nucleosome near SHL +2 and causes structural rearrangements

To dissect the mechanism of nucleosome invasion by FoxO1, we expressed a truncated human FoxO1 construct (residues 144–266) encompassing the DNA-binding domain flanked by short N- and C-terminal regions (Figure 1e, Figure S2a). We first assessed FoxO1 binding to IGFBP1 nucleosomes by electrophoretic mobility shift assays (EMSA). At nucleosome:FoxO1 molar ratios between 1:10 and 1:20, with the nucleosome concentration held at 300 nM, the entire nucleosome population was shifted, indicating quantitative binding of FoxO1 under these conditions (Figure 1f), which were subsequently used to prepare IGFBP1 nucleosome–FoxO1 complexes for cryo-EM characterization.

Our structural analysis revealed a heterogeneous population that included free IGFBP1 nucleosomes, stacked nucleosomes, and a subset of nucleosomes containing an additional bound density that we attributed to FoxO1 (Figure S3). After iterative rounds of classification of the FoxO1-bound subset, we determined the structure of the IGFBP1 nucleosome–FoxO1 complex (^IGFBP1^Nuc-FoxO1) at an overall resolution of 6.2 Å (Figure 1g, Figure S8 and Table S1).

The structure shows density with the size expected for the FoxO1 construct, positioned on the nucleosomal DNA near SHL +2, consistent with the location of the insulin-responsive element (IRE) site mapped in the FoxO1-free IGFBP1 nucleosome Class I (Figure 1g and Figure 1a). Also, we observed that although FoxO1 binds to this internal nucleosomal position, its association induces an increase in distal DNA unwrapping from 8 to ∼15 bp, with the final histone–DNA contact occurring near SHL +5.5 while the proximal DNA arm remains unaltered (Figure 1g).

In addition to the effect on the distal DNA arm, binding of FoxO1 to the IGFBP1 nucleosome induces allosteric destabilization of the histone octamer core, manifested as local unfolding of secondary structural elements in the H2A C-terminal and H3 N-terminal regions (located distal to the FoxO1 binding site) (Figure 1h, left), and the repositioning of the H2A–H2B histone dimer adjacent to the unwrapped distal DNA (Figure 1h, right). This destabilization also contributes to the conformational heterogeneity and lower resolution of the reconstruction.

### HNF1α induces outer unwrapping of the IGFBP1 Nucleosome

To elucidate the molecular mechanism by which HNF1α engages its target DNA motif within the IGFBP1 nucleosome, we expressed a truncated human HNF1α construct (residues 90–288) encompassing the POU-specific (POU_S_) domain and POU-homeodomain (POU_H_), flanked by short unstructured N- and C-terminal regions (Figure 2a, Figure S2b). Our binding assays showed partial shifting of the IGFBP1 nucleosome population at a nucleosome:HNF1α molar ratio of 1:5, with complete shifting observed at a ratio of 1:10 (nucleosome concentration maintained at 300 nM) (Figure 2b). The latter condition was used to prepare IGFBP1 nucleosome–HNF1α complexes (^IGFBP1^Nuc-HNF1α) for cryo-EM study.

**Figure 2:**
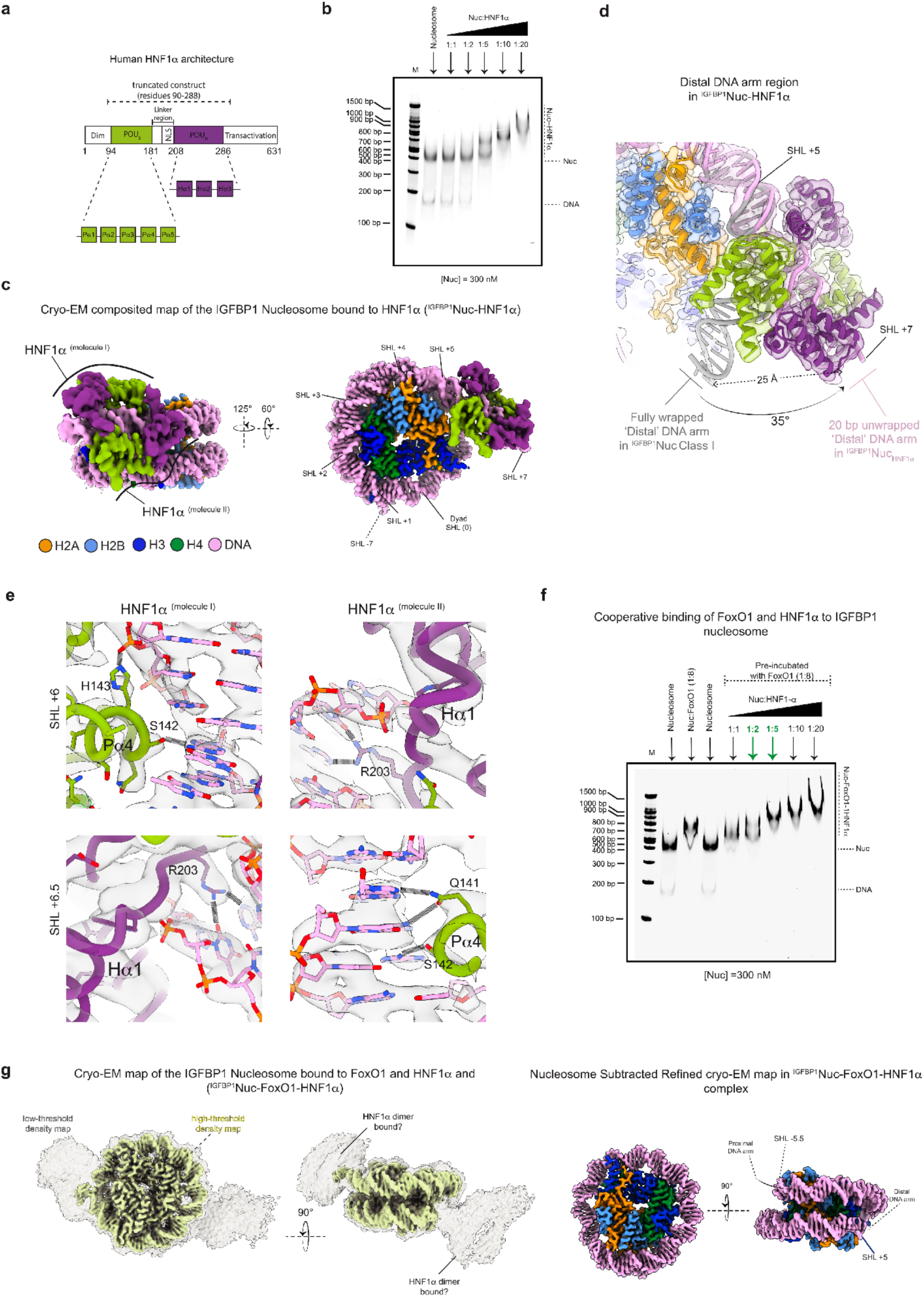
Cryo-EM structure of the *IGFBP1* Nucleosome bound to HNF1α (^IGFBP1^Nuc–HNF1α). **a.** Schematic of the human HNF1α domain architecture highlighting the POU-specific (POU_S_) domain and POU-homeodomain (POU_H_). The truncated construct used in this work (residues 90-288) is indicated. **b.** Representative EMSA (5% TBE gel) showing binding of HNF1α to *IGFBP1* nucleosomes. The nucleosome:HNF1α molar ratios used are indicated. **c.** Cryo-EM composite map of *^IGFBP1^*Nuc–HNF1α in different views. Each molecule of the HNF1α dimer bound to the distal DNA is labeled and colored by domain as in (a). Color key for the nucleosome is indicated and relevant SHLs are annotated. **d.** Close-up view of the unwrapped distal DNA arm in *^IGFBP1^*Nuc–HNF1α relative to the fully wrapped *^IGFBP1^*Nuc_Free_ Class I structure (gray ribbon). **e.** Interactions between the HNF1α dimer and the distal nucleosomal DNA. The density map is shown as a transparent grey surface with the atomic model fitted. Residues establishing hydrogen bonds are labeled. **f.** Representative EMSA (5% TBE gel) showing simultaneous binding of FoxO1 and HNF1α to *IGFBP1* nucleosomes. The nucleosome:HNF1α molar ratios are indicated. Nucleosomes were pre-incubated with FoxO1 at 1:8 (nucleosome:FoxO1) molar ratio. Green arrows mark early shifting of nucleosomes by HNF1α in the presence of FoxO1 compared to HNF1α alone (panel b). **g.** Cryo-EM density map of the ^IGFBP1^Nuc–FoxO1–HFN1α complex. Lef: High-threshold (yellow solid density) and low-threshold/low-pass filtered (gray transparent surface) representation of the cryo-EM map. The nucleosome is unwrapped from both DNA ends. Right: Colored high-threshold cryo-EM ^IGFBP1^Nuc–FoxO1–HFN1α map. with DNA register indicated. The proximal and distal DNA arms appear unwrapped by about 15 bp and 20 bp, respectively.

In the cryo-EM composite map of the ^IGFBP1^Nuc-HNF1α, the nucleosome core and the HNF1α–DNA interacting regions were solved at 3.1 Å and 3.9 Å overall resolution, respectively (Figure S4, Figure S8 and Table S1). Two HNF1α molecules (see molecule I and II in Figure 2c) are well-resolved and bind as a dimer contacting both DNA grooves spanning SHLs +5.5 to +6.5, consistent with the predicted position of the binding site inferred from the IGFBP1 nucleosome Class I structure (Figure 2c and Figure 1a). The ^IGFBP1^Nuc-HNF1α displays a pronounced unwrapping of approximately 20 bp of the distal DNA arm, with the last DNA-histone contact observed at SHL +5 (Figure 2d). The unwrapped DNA is straightened (unbent) by ∼35° and displaced ∼25 Å from the histone octamer core, adopting a fully extended conformation stabilized by the HNF1α dimer, which engages the DNA in a pseudo-symmetric arrangement (Figure 2d).

HNF1α^molecule I^ contacts the DNA major groove at SHL +6 via residues S142 and H143, located in the Pα4 helix of the POU_s_ domain, establishing hydrogen-bond interactions with DNA bases and the phosphate backbone, respectively (Figure 2e, left). In contrast, HNF1α^molecule II^ engages the DNA minor groove at the same location through residue R203, located in the N-terminal loop adjacent to helix Hα1 of the POU_H_ homeodomain, which hydrogen-bonds with two DNA bases (Figure 2e, right). At SHL +6.5, residue R203 of the HNF1α^molecule I^ contacts the DNA minor groove, while residues Q141 and S142 within the POUs domain of HNF1α^molecule II^ make hydrogen-bonds and polar interactions with DNA bases in the major groove. Overall, the HNF1α-DNA interacting region in our complex, resembles the previous X-ray structure reported for HNF1α bound to a linear DNA fragment (PDB 1IC8)^22^. Superimposing the nucleosomal and linear DNA structures, using the DNA binding motif as a reference, reveals that both HNF1α monomers on the nucleosome appear slightly rotated toward the histone octamer core relative to their position when bound to linear DNA (Figure S7).

Together, our structures reveal distinct modes of nucleosome engagement by HNF1α and FoxO1. While FoxO1 binds an internal superhelical location, destabilizing the nucleosome and enhancing distal DNA unwrapping, HNF1α binds near the DNA entry–exit region, stabilizing a persistently unwrapped conformation. These observations point to a potential mechanism by which FoxO1 and HNF1α may affect each other’s binding and suggest that they can act synergistically on the same nucleosome to modulate its structure and stability.

### FoxO1 facilitates HNF1α binding and promotes extensive nucleosome unwrapping

We next examined whether FoxO1 and HNF1α can bind cooperatively to the same IGFBP1 nucleosome and how their combined action remodels nucleosome architecture. EMSAs in which IGFBP1 nucleosomes were pre-incubated with FoxO1 at varying molar ratios prior to the addition of HNF1α showed a positive effect of FoXO1 on HNF1α nucleosome binding (Figure S9). With the nucleosome concentration maintained at 300 nM, pre-incubation with FoxO1 at a 1:8 nucleosome:FoxO1 molar ratio enabled partial and complete shifting of the nucleosome population upon addition of HNF1α at 1:2 and 1:5 nucleosome:HNF1α molar ratio, respectively (Figure 2f). By comparison, HNF1α alone required a molar ratio of 1:10 nucleosome:HNF1α to achieve full nucleosome binding under our binding conditions (Figure 2b). These results show that FoxO1 facilitates the engagement of HNF1α with the IGFBP1 nucleosome by approximately two-fold and suggest binding cooperativity.

To investigate the possibility that this cooperative binding could promote larger-scale rearrangements of nucleosome architecture, we assembled IGFBP1 nucleosome–FoxO1–HNF1α ternary complexes under the binding conditions defined above and subjected them to cryo-EM analysis. This sample appeared highly heterogeneous, with nucleosomes exhibiting a range of DNA unwrapping states. Out of this ensemble, the most stable class reached 3.5 Å overall resolution (^IGFBP1^Nuc-FoxO1-HNF1α) and showed a well-resolved nucleosome core particle at high density thresholds (Figure S5, Figure S6 and Table S1). At lower thresholds, two additional densities emerge at opposite sides of the nucleosome, indicating pronounced unwrapping at both DNA entry–exit sites (Figure 2g, left). Although the conformational flexibility of these peripheral densities precluded high-resolution modeling, their size and position are consistent with the binding of a HNF1α dimer at each nucleosomal DNA arm. Focused refinement of the nucleosome core yielded a map at 3.1 Å resolution, enabling unambiguous assignment of the DNA register and revealing asymmetric DNA unwrapping. The distal DNA arm unwraps by approximately 20 bp, with the final histone–DNA contact near SHL +5, whereas the proximal DNA arm unwraps by ∼15 bp, with its final contact near SHL −5.5 (Figure 2g, right). These results demonstrate that the destabilization of the nucleosome caused by pre-binding of FoxO1 enables HNF1α to bind and unwrap the DNA on both sides of the nucleosome, a behavior not observed for the IGFBP1 nucleosome in the presence of either TF individually.

### Mechanical unwrapping of the free IGFBP1 nucleosome

To gain insight into how these TFs modulate nucleosome mechanical stability to enable access to their DNA target motifs, we used high-resolution dual-trap optical tweezers to mechanically induce nucleosome unwrapping in the absence or presence of FoxO1 or/and HNF1α (Figure 3a). To facilitate comparisons across single-molecule measurements, all analyses presented here are based on the first pulling trajectories of intact nucleosomes, which are identified when the Force (pN) vs Extension (nm) curves exhibit both a low-force and a high-force transition, as previously described for the Widom 601 sequence^32^. The low-force transition corresponds to the unwrapping of about 27 bp from one of the entry/exit arms of the nucleosome, whereas the high-force transition corresponds to the unwrapping of the rest of the DNA around the histone core.

**Figure 3:**
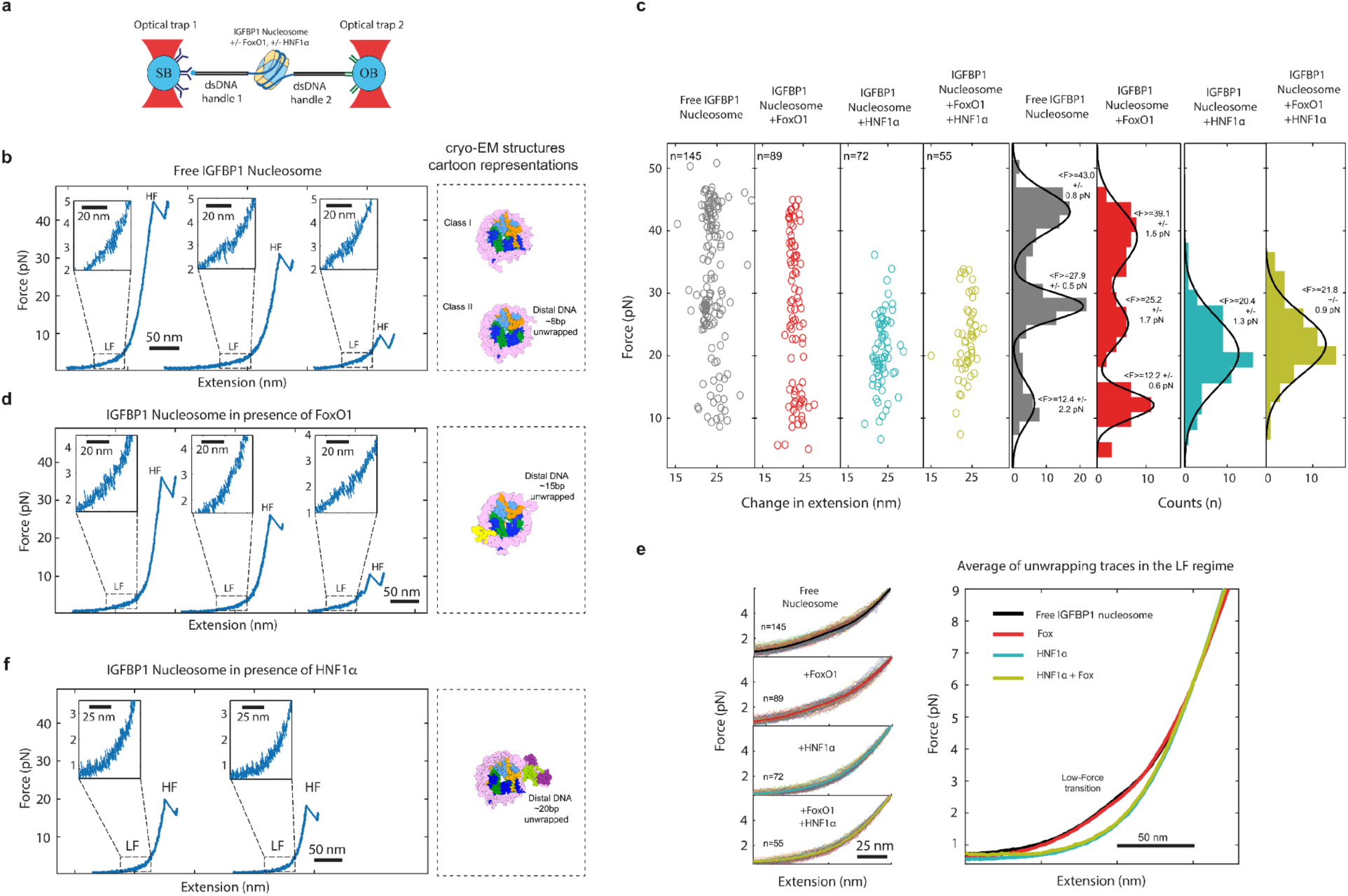
Mechanical unwrapping of IGFBP1 Nucleosomes and the effect of FoxO1 and HNF1α. **a.** Dual-trap high-resolution optical tweezers experimental set-up. The nucleosome was ligated to DNA handles, which were then tethered to 1 μm polystyrene microbeads by ligation to DNA oligo coated beads and by biotin binding to streptavidin coated beads. **b.** Representative Force (pN) vs Extension (nm) unwrapping traces of the free *IGFBP1* nucleosome, FoxO1–bound and HNF1α–bound complexes (left), respectively. Cartoon representation of their corresponding cryo-EM structure is shown (right). **c.** Plot of the Force (pN) vs Extension (nm) datapoints of the individual high-force ‘rip’ transitions obtained for the free *IGFBP1* nucleosome, FoxO1–bound (+FoxO1), HNF1α–bound (+HNF1α) and both factors bound (+FoxO1, +HNF1α) conditions. For the free and +FoxO1 nucleosomes, the data points were adjusted to a tri-modal distribution (R^2^=0.9). For the +HNF1α and for the +FoxO1+HNF1α cases, the datapoints were adjusted to a unimodal distribution (R^2^=0.9) (see Methods). **d.** Representative Force (pN) vs Extension (nm) unwrapping traces of the *IGFBP1* nucleosome bound to FoxO1 (left). Cartoon representation of its corresponding cryo-EM structures is shown (right). **e.** Left: Multiple individual unwrapping traces (thin lines) and their average (thick line) for the free *IGFBP1* nucleosome, +FoxO1, +HNF1α, and both factors bound (+FoxO1, +HNF1α) conditions. Right: Overlapping of the average traces obtained per condition and compared at the low-force regime. **f.** Representative Force (pN) vs Extension (nm) unwrapping traces of the *IGFBP1* nucleosome bound to HNF1α (left). Cartoon representation of its corresponding cryo-EM structures is shown (right).

For the free IGFBP1 nucleosomes, we observed that the low-force transition, associated with outer DNA unwrapping, occurs at a force of about 2–3 pN and displays a non-cooperative behavior, and a barely detectable change in extension (Figure 3b). These features differ from those of the canonical Widom 601 nucleosomes, for which the low-force transition occurs at ∼4–5 pN and takes place either in a non-cooperative or a cooperative manner, with a change in extension of ∼11 nm or ∼20 nm, respectively^32^. These results indicate that nucleosomes assembled on the native IGFBP1 DNA exhibit an increased propensity for spontaneous unwrapping (with a low-force transition extension significantly lower than 11 nm), consistent with enhanced *nucleosome breathing*. This interpretation agrees with our structural analysis of the free IGFBP1 nucleosome, in which the Class II conformation reflects spontaneous unwrapping of approximately 8 bp of the distal DNA arm. Notably, as mentioned above, this region of the IGFBP1 sequence is GC rich and exhibits a reduced intrinsic bendability and a higher tendency to unwrap spontaneously, consistent with previous single-molecule studies on 601 nucleosomes^35,33^.

In the high-force regime of the force-extension traces associated with inner DNA unwrapping, the free IGFBP1 nucleosome exhibits a cooperative transition or ‘rip’ (Figure 3b), characterized by a broad distribution of forces peaked around 43.0 ± 0.8 pN, 27.9 ± 0.5 pN, and 12.4 ± 2.2 pN (Figure 3c). Interestingly, a similar “tri-modal” distribution of high-force transition has previously been reported for canonical Widom 601 nucleosomes, indicating an intrinsic structural plasticity of the nucleosomes that is independent of the DNA sequence^32^.

### FoxO1 destabilizes the IGFBP1 nucleosome

We next performed mechanical unwrapping experiments on individual IGFBP1 nucleosomes in the presence of 0.5 µM FoxO1. The force–extension traces revealed a weak, difficult-to-discern non-cooperative low-force transition, occurring at an average force of ∼2–3 pN (Figure 3d). To quantitatively assess the effect of FoxO1 on outer DNA unwrapping, we averaged multiple first pulling force–extension traces and compared them in their low-force regime (Figure 3e, left). We find that the average behavior of the IGFBP1 nucleosome unwrapping trajectories is not significantly affected at the low-force regime by FoxO1 binding (Figure 3e, right). On the other hand, in the high-force regime, the IGFBP1 nucleosomes in the presence of FoxO1 continued to display a cooperative rip transition with a tri-modal distribution, with force peaks centered at 39.1 ± 1.5 pN, 25.2 ± 1.7 pN, and 12.2 ± 0.6 pN (Figure 3c). These values are uniformly reduced relative to those obtained for the free nucleosomes, with the lower peak centered at ∼12 pN being more pronounced, indicating a higher population of this less stable “mechanical conformation” in the presence of FoxO1. Together, our results indicate that although FoxO1 does not cause a significant mechanical effect on outer DNA wrapping of the IGFBP1 nucleosome, binding of this factor to its cognate sites causes a global destabilization of the DNA-histone core interactions. These results are consistent with our FoxO1-bound cryo-EM structure, in which FoxO1 binding is associated with enhanced distal DNA unwrapping accompanied by allosteric destabilization of the histone octamer core (Figure 1g and 1h).

### HNF1α affects IGFBP1 nucleosome stability and reshapes its unwrapping trajectory

In the presence of HNF1α, IGFBP1 nucleosomes display a nearly undetectable low-force unwrapping transition (Figure 3f), indicating that histone–DNA contacts at one of the DNA entry–exit sites are substantially perturbed. This observation is supported by previous mechanical unwrapping studies of Widom *601* nucleosomes combined with smFRET detection, which revealed that the stiffer (higher GC content) DNA arm is the first to unwrap during the low-force transition^32,33^. These observations are consistent with the ^IGFBP1^Nuc–HNF1α cryo-EM structure, in which two HNF1α molecules unbend and maintain approximately 20 bp of the distal DNA arm between SHLs +5 and +7 in an unwrapped state. The average of multiple unwrapping traces at this low-force regime (Figure 3e) further indicates that binding of HNF1α results in a pronounced suppression of the low-force transition on the IGFBP1 nucleosome (in contrast to the effect of FoxO1), consistent with its direct engagement with the distal DNA region and its large, induced unwrapping.

In the high-force regime, the HNF1α-bound IGFBP1 nucleosomes exhibit a single, well-defined force population centered at 20.4 ± 1.3 pN. This behavior is significantly distinct from the tri-modal distribution observed for the high-force transition of apo IGFBP1 nucleosomes and those bound to FoxO1.

Notably, mechanical unwrapping experiments performed in the presence of both FoxO1 and HNF1α yielded force–extension signatures indistinguishable from those observed in the presence of HNF1α alone (Figures 3c and 3e). These results indicate that, although FoxO1 may facilitate the recruitment of HNF1α to the nucleosome, HNF1α binding ultimately dominates the mechanical response, both reducing and constraining nucleosome stability along a single disassembly pathway.

### FoxO1 and HNF1α modulate the IGFBP1 nucleosome disassembly dynamics

We next asked whether the TF–dependent changes in IGFBP1 nucleosome mechanics described here affect the stability of the nucleosome in successive pulling–relaxation cycles. Previous correlated optical tweezers and single-molecule FRET studies have established characteristic mechanical signatures for the unwrapping of canonical Widom 601 nucleosomes and their subnucleosomal derivatives, including hexasomes and tetrasomes^32^. In addition, atomic force microscopy studies have shown that nucleosomes can spontaneously disassemble in solution through the release of H2A–H2B dimers^31^. Together, these observations indicate that nucleosomes can access multiple disassembly pathways, whose relative probabilities can be quantified using single-molecule force spectroscopy.

Accordingly, we investigated how FoxO1 and HNF1α bias nucleosome integrity and fate under mechanical stress by analyzing the second pulling–relaxation cycle. Starting from an intact IGFBP1 nucleosome (first pulling, “octasome”), the next force–extension-followed-by-relaxation traces revealed the presence of hexasomes, tetrasomes, or bare DNA, which were assigned based on their previously established mechanical signatures features^32^(see Methods) (Figure 4a–c).

**Figure 4:**
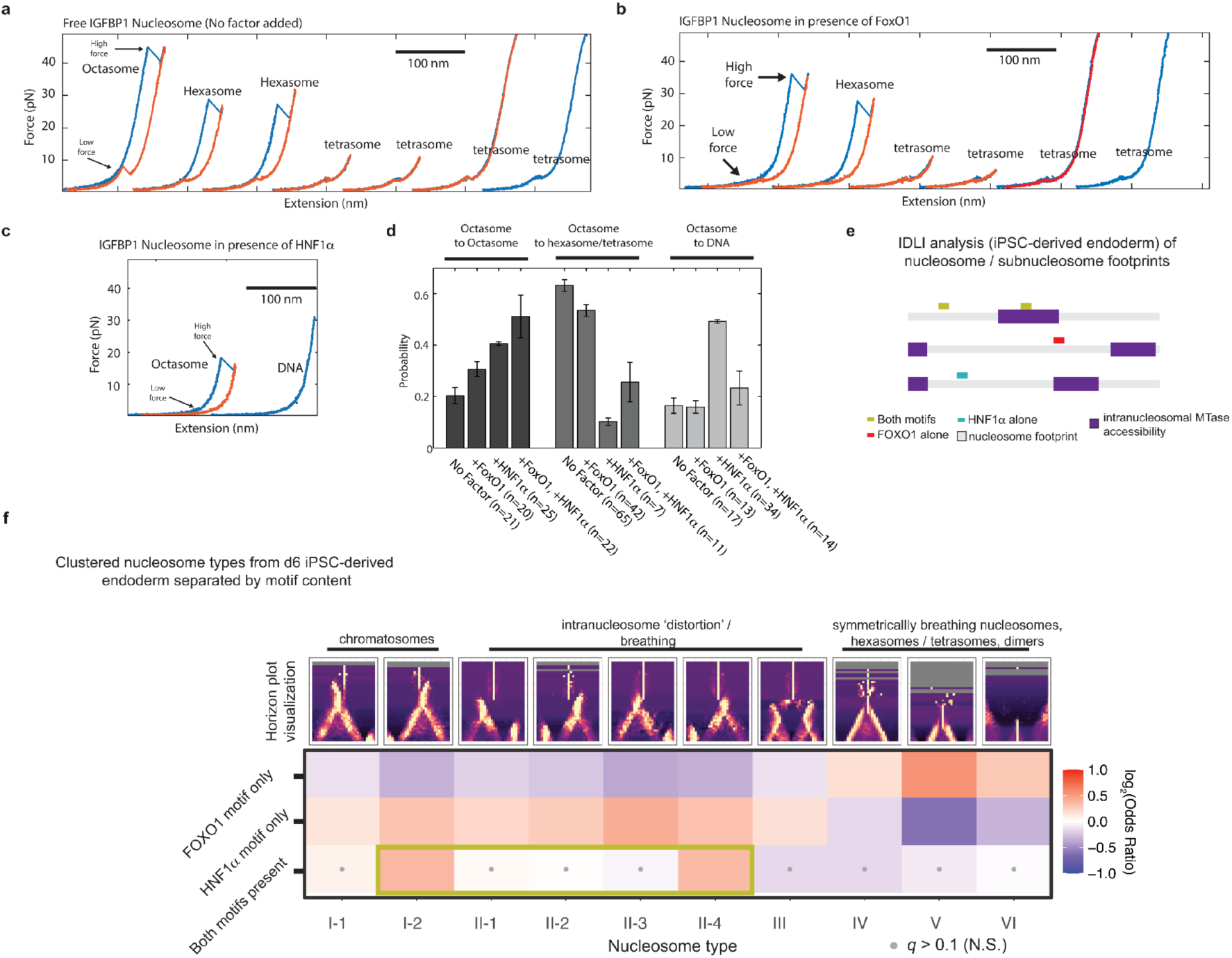
Analysis of the IGFBP1 Nucleosome disassembly, subnucleosome-resolution footprinting and the effect of FoxO1 and HNF1α. **a-c.** Representative traces of multiple pulling-relaxation cycles (blue–orange) for the free *IGFBP1* nucleosome, and in presence of FoxO1 or HNF1α, respectively. These successive traces show the disassembly of nucleosome into hexasomes, tetrasomes, and naked DNA (as indicated). **d.** Bar graph showing the probabilities of IGFBP1 nucleosome disassembly into hexasomes, tetrasomes, or naked DNA across different experimental conditions. The number of datapoints collected per condition is indicated by ‘n’. **e.** Schematic of the IDLI pipeline and its application to this study. Nucleosome footprints called in Yang et al.^36^ and analyzed using the IDLI pipeline were filtered to define nucleosomes within hepatocyte-active regulatory regions containing FoxO1and/or HNF1α binding motifs. The IDLI nucleosome clustering pipeline was then used to examine nucleosome or subnucleosome abundance conditioned on motif presence within each called footprint. **f.** Enrichment analyses for nucleosome types defined at these regulatory loci using the IDLI pipeline. Red heat indicates enrichment of specific nucleosome types with respect to the background distribution of all nucleosome calls, while blue indicates concomitant depletion. All effect sizes are visualized as log-transformed effect sizes derived from Fisher’s exact tests, which are then multiple-hypothesis-test corrected using Storey’s q value. All tests that are not significant (N.S.) below a False Discovery Rate (FDR) threshold of 10% are marked with a grey dot.

Analysis of the apo IGFBP1 nucleosomes showed that ∼20% of molecules remained as intact octasomes in the second pulling cycle, whereas ∼63% transitioned to subnucleosomal states (hexasomes or tetrasomes). Direct conversion from octasomes to bare DNA was comparatively less probable (∼17%) (Figure 4d). The presence of FoxO1 did not significantly alter these disassembly probabilities, indicating a limited impact on the natural ‘plasticity’ behavior of the IGFBP1 nucleosome, as we saw reflected on their fate under repeated mechanical perturbation (Figure 4d).

In contrast, in the presence of HNF1α the probability of retaining an intact octasome increased to ∼41%, whereas the combined probability of disassembling into hexasomes or tetrasomes decreased to ∼10%, substantially lower than that observed for apo nucleosomes (∼63%) or those bound to FoxO1 (∼50%) (Figure 4d). On the other hand, HNF1α increased the probability of direct octasome-to-bare-DNA transitions to ∼49% (Figure 4d), an all or nothing behavior

Interestingly, in the presence of both FoxO1 and HNF1α, IGFBP1 nucleosomes exhibited the highest probability of remaining intact as octasomes (∼51%) in the second pulling–relaxation cycle. Under these conditions, intermediate probabilities were observed for disassembly into hexasomes or tetrasomes (∼26%) and for direct transitions to bare DNA (∼23%) (Figure 4d), indicating a combined influence of both TFs on nucleosome fate that could not have been predicted from their individual effects.

### FoxO1 and HNF1α binding to cognate DNA motifs in promoter regions modulate nucleosomes architecture and stability

To explore how the presence of FoxO1 and/or HNF1α binding motifs may influence the stability of nucleosomes in cells, we re-analyzed published subnucleosome resolution adenine methyltransferase single-molecule footprinting data (the ‘SAMOSA’ assay^34^) from human induced pluripotent stem cells differentiated into endoderm (Figure 4e)^35^. Leveraging the Iteratively Defined Lengths of Inaccessibility (IDLI) pipeline for systematically identifying chromatosomes, distorted nucleosomes, and subnucleosomes, we focused our analysis on DNase I hypersensitive sites (DHS) profiled in human primary hepatocytes^35^, which are the most likely regions of the genome to play roles in regulating transcription. Since TFs must recognize their cognate DNA sequence motifs in chromatinized environments, we can safely assume that if these motifs are present within the DHS, they are bona fide binding sites for these (or closely related) TFs^36^. First, we computationally identified nucleosome footprints falling within DHS, then filtered these nucleosome footprints for those that contain a high-confidence match to the FoxO1 motif, HNF1α motif, or both. Our IDLI nucleosome clustering defined 10 distinct classes of (sub)nucleosomes (*i.e.* nucleosome ‘types’) (Figure 4f, top), which fell into six broad groups as defined by hierarchical clustering (Figure S10a). Nucleosome types broadly matched those we previously identified and biochemically/genetically validated^37^, including chromatosomes, distorted nucleosomes with focal methyltransferase accessibility at regions varying from SHL 0/±1 to SHL ±6, breathable nucleosomes, and subnucleosomes (Figure S10b).

To examine how the presence of specific TF motifs within these identified footprints influences nucleosomes’ architecture, we assessed the statistical enrichment (red) or depletion (blue) of specific nucleosome types for footprints containing either FoxO1, HNF1α, or both motifs, which we visualized as a heatmap of log-transformed effect sizes (Figure 4f, bottom). This analysis revealed that the presence, absence, or combination of these motifs had significant effects on the wrapping state of nucleosomes, their accessibility, and the presence of subnucleosome species. For regions harboring the FoxO1 binding motif alone, we observed the highest representation of unwrapped nucleosomes and subnucleosomal species, such as types IV, V and VI (hexasomes/tetrasomes, H2A/H2B dimers and TF footprints, respectively). For regions containing only the HNF1α motif, the largest populations were those of stably wrapped nucleosomes and of ‘breathing’ nucleosomes with accessible intra-nucleosome DNA (*e.g.* I-1, II-1 to II-4), with a reduced population of subnucleosome species (types IV to VI) compared to footprinting regions harboring the FoxO1 motif alone. Interestingly, regions containing both FoxO1 and HNF1α binding motifs display an intermediate phenotype, with a larger presence of stably wrapped nucleosomes and ‘breathing’ nucleosomes species relative to sites with the FoxO1 motif alone, but smaller relative to sites with only the HNF1α cognate motif (Fig. 4g). This intermediate phenotype also extended to the representation of footprints consistent with subnucleosomes (*e.g.* types IV, V and VI; likely hexasomes/tetrasomes), with HNF1α alone having the fewest subnucleosomes, and FoxO1 alone the greatest. These results suggest that the combined presence of these lineage-specific TFs directly modulates the stability of specific nucleosomal species in cells.

Our *in vivo* observations are in general agreement with our cryo-EM and optical tweezer findings, even when the *in vivo* regulatory environment is significantly more complex with respect to possible nucleosome-interacting factors (*e.g.* remodelers, HMG-family proteins, linker histone) and nucleosomal sequences (all sequences enriched in hepatocyte DHS, versus a single *IGFBP1* sequence). Still, nucleosomes with motifs for HNF1α alone, FoxO1 alone, and the combination of the two, each show distinct structural features that are qualitatively consistent with the structural and biophysical data.

IGFBP1 nucleosomes in the presence of FoxO1 alone show the most unwrapped architecture, a perturbed histone octamer core, and exhibited a reduced stability during mechanical unwrapping, with a high tendency (∼50%) to disassemble into subnucleosome species (hexasome/tetrasome).These structural and biophysical results align well with our *in vivo* footprinting observation that nucleosomes harboring a FoxO1 motif alone, are the most enriched in subnucleosomal particles and the most accessible nucleosomes. HNF1α−bound IGFBP1 nucleosomes, while unwrapped, display an unaltered histone octamer core, and these nucleosomes demonstrate the lowest disassembly probability (∼10%), again in agreement with our *in vivo* footprinting results showing that nucleosomes with the HNF1a-motif alone are the least methyltransferase accessible and the most enriched for intact nucleosomes. In the presence of both FoxO1 and HNF1a, we observe an IGFBP1 nucleosome structure that is unwrapped from both sides yet retains an unperturbed histone octamer core. These nucleosomes have an intermediate probability of subnucleosome disassembly (∼26%) and, through successive pulling/relaxing cycles, have the highest probability (∼51%) of remaining as octasomes. These results are also consistent with our footprints data showing an intermediate distribution of wrapped, unwrapped, and subnucleosomal states among FoxO1. Taken together, these results provide *in vivo* evidence supporting the notion that HNF1a and FoxO1 can regulate the stability and wrapping state of nucleosomes.

## DISCUSSION

Our work shows that nucleosomes assembled on a native IGFBP1 promoter sequence display both fully wrapped and partially unwrapped states, where the DNA arm enriched in GC content spontaneously detaches from the histone octamer core, resulting in significantly smaller low-force transition under force than the strongly wrapped Widom 601 nucleosome. Thus, sequence-dependent mechanical properties of endogenous DNA drive asymmetric unwrapping and spontaneous nucleosome breathing, in agreement with previous studies showing that nucleosomes can display spontaneous opening, with the stiffer DNA arm having a preference to unwrap first ^31,38,39^. This mechanical asymmetry of the nucleosomal DNA has been found throughout the genome, suggesting that it plays a significant role in the control of gene expression^5^.

We found that FoxO1 engages the IGFBP1 nucleosome at an internal superhelical location and enhances distal DNA unwrapping. This unwrapping is accompanied by the re-positioning of the adjacent H2A–H2B histone dimer and the local unfolding of its neighbor H3 N-terminal and the H2A C-terminal regions. Thus, FoxO1 destabilizes the histone octamer core by an allosteric mechanism likely mediated through the nucleosomal DNA. Previous studies have reported that other pioneer factors, such as SOX11, are also able to destabilize the nucleosomal DNA arm distal from the factor binding site^10^ and that the distal H2A–H2B histone dimer can reposition upon DNA unwrapping^40^. This ‘allosteric destabilization’ model is supported by our optical-tweezers results, which show that FoxO1 reduces the global mechanical stability of the IGFBP1 nucleosome, facilitating inner DNA unwrapping while preserving the intrinsic ‘plasticity’ of the nucleosome to sample different molecular trajectories during its mechanical unraveling.

Our pulling–relaxation cycles of free IGFBP1 nucleosomes reveal that they frequently transition into subnucleosomal intermediates, rather than remaining intact as octasomes or fully disassembling into bare DNA. Notably, FoxO1 binding does not significantly alter these disassembly probabilities, suggesting that by enhancing the distal unwrapping and destabilizing the histone core, as observed in our cryo-EM results, this pioneer factor preserves the ‘plasticity’ of the IGFBP1 nucleosome. A direct destabilizing role for FoxO1 on the nucleosome, is supported by the reanalysis of *in vivo* methyltransferase footprinting data in human iPSC-derived endoderm^36^. These findings showed that nucleosomes within regulatory regions containing a FoxO1 motif are enriched for unwrapped nucleosomes and subnucleosomal species compared to nucleosomes with a HNF1α motif, or nucleosomes containing both HNF1α and FoxO1 motifs.

On the other hand, our IGFBP1 nucleosome–HNF1α structure shows how binding of two HNF1α molecules to the target motif results in an extensive unwrapping of the DNA distal arm, which is stabilized in an extended conformation by multiple contacts with the POU_S_ and POU_H_ domains. Although previous crystallography studies showed that HNF1α binds to its target motif on linear DNA^23,26^ in a similar fashion, its binding implications in a nucleosomal context were unknown. Moreover, other TFs also containing POU_S_ and POU_H_ domains, like OCT4, exhibit different modes of nucleosome engagement depending on the location of their target motif. For instance, when in cooperation with SOX2, OCT4 binds nucleosomal DNA at superhelical location ±6 and only its POU_S_ domain is involved in the interaction, whereas both POU_S_ and POU_H_ domains are engaged when OCT4 binds to linker DNA outside the nucleosome core, resulting in distinct effect on the nucleosome^41,42^. Together, these observations indicate that TF binding mode and the resulting remodeling outcome are strongly dependent on the position of the target motif within the nucleosome.

Our findings show that HNF1α engages the nucleosome very differently from FoxO1, causing large distal DNA unwrapping while not disturbing the histone octamer, and constraining the inner DNA unwrapping to follow a unimodal high-force transition. In fact, HNF1α appears to increase the stability of the histone core, which either remains intact as an octasome or undergoes full ejection from the DNA in successive pulling-relaxation cycles. These results suggest that, by unwrapping the outer DNA, HNF1α reduces the generation of incremental structural changes that would otherwise arise from repeated disruptions of DNA-histone contacts, leaving the histone core undisturbed. We propose that by increasing the rigidity of the histone core and by promoting the unwrapping of the outer DNA, HNF1α prevents the spontaneous release of H2A-H2B histone dimers, favoring a limited set of disassembly outcomes.

These in vitro results agree with our nucleosome footprinting findings in iPSC-derived human endoderm cells showing that the presence of HNF1α only motifs result in a shift towards more stable nucleosomes compared to nucleosomes containing FoxO1 only motifs, or a combination of both motifs. Of note, the notion of a TF stabilizing nucleosome structures is not without precedent. We previously demonstrated that in similar footprinting experiments in mouse embryonic stem cells (mESCs), rapid protein degradation of SOX2 at SOX2-binding sites resulted in de-stabilization of local nucleosomes, suggesting that SOX2 itself may contribute to the stability of core nucleosomes within cells^36^.

Additionally, our in vitro assays show that FoxO1 enhances binding of HNF1α to the IGFBP1 nucleosome, while force-induced nucleosome unwrapping in the presence of both FoxO1 and HNF1α is indistinguishable from that observed with HNF1α alone. These results suggest that FoxO1 facilitates nucleosome HNF1α recruitment, and HNF1α controls the nucleosome mechanical response. In the presence of both factors, nucleosomes exhibit the highest reported probability to remain as intact octasomes but still display alternative disassembly pathways into hexasomes/tetrasomes or bare DNA. We propose that by pre-existing in an “open” state, nucleosomes bound by these factors become more amenable to unwrapping under mechanical disruption by an external force. Continual mechanically induced unwrapping leads the nucleosome to adopt a range of conformations under tension, which may create opportunities for H2A–H2B dimer ejection leading to the formation of hexasomes or tetrasomes. These in vitro results are qualitatively consistent with the distribution of distinct nucleosome types we observe at sites containing FoxO1 and HNF1α motifs, which are skewed towards subnucleosomes more than those with only HNF1α motifs, but more towards stably wrapped species compared to with sites with only a FoxO1 motif.

The general agreement of our in vitro and in-cell measurements is notable, given that *in vivo* nucleosomes are also substrates for a wide range of other chromatin-associated activities. Even in the complex regulatory environment of a cell, our data show that nucleosomes capable of accommodating both FoxO1 and HNF1α binding are structurally distinct from nucleosomes containing either TF motif alone. ATP-dependent chromatin remodeling enzymes exert mechanical stress on nucleosomes, resulting in nucleosome/subnucleosome sliding or disassembly. Our structural and biophysical data suggest that TFs can (individually or in combination) directly alter the wrapping state of nucleosomes in a programmed fashion, in the absence of an ATP-dependent chromatin remodeling enzyme. In cells, however, TFs directly engage chromatin remodeling enzymes^43^, possibly via specific structural motifs^44^. Our *in vivo* observations likely result from the combined activities of the TFs themselves and of ATP-dependent remodelers, and other chromatin regulatory factors (*e.g.* chaperones, acetyltransferases, etc.).

Altogether, our results support a model wherein TFs quantitatively modulate the structural properties of nucleosomes, providing much-needed structural and biophysical context to the ‘pioneer factor’ model. Given the combinatorial complexity of both TFs and their associated binding sites within eukaryotic chromatin, our results highlight the need for more nucleosome-bound TF structures on physiological DNA sequences. Future work may focus on discerning the extent to which different TFs remodel nucleosome structure on such DNA templates, and the possible stimulatory or antagonistic roles for chromatin co-factors.

## METHODS

### Histone Purification and Octamer assembly

Xenopus laevis histones for nucleosome assembly were overexpressed in the Escherichia coli BL21(DE3) strain and purified from inclusion bodies as previously described^1^. Briefly, *Escherichia coli* was grown at 37 °C and induced with 0.2 mM IPTG for 3 h. Cells were harvested, resuspended in lysis buffer (100 mM NaCl, 20 mM HEPES pH 7.5, 0.05% Triton X-100), and lysed by sonication. The lysate was centrifuged at 17,000 × g for 30 min at 4 °C, and the insoluble pellet containing inclusion bodies was washed repeatedly with lysis buffer until white. The pellet was resuspended in unfolding buffer (6 M guanidine hydrochloride, 20 mM HEPES pH 7.5, 5 mM DTT) and incubated for 5h at 4 °C, followed by centrifugation at 17,000 × g for 30 min to remove insoluble material. The supernatant was dialyzed against buffer containing 7 M urea, 100 mM NaCl, 20 mM sodium acetate pH 5.2, and 5 mM DTT, and individual histones were purified by cation-exchange chromatography on a HiTrap SP column using a linear NaCl gradient from 100 mM to 500 mM in 7 M urea, 20 mM Sodium acetate pH 5.2, and 5 mM DTT. Fractions were analyzed by SDS–PAGE, pooled, and dialyzed into buffer containing 5 mM DTT for further use. To assemble the octamer, the four histones were mixed at an equimolar ratio in unfolding buffer containing 6 M guanidine hydrochloride, 20 mM Tris–HCl (pH 7.5), and 5 mM DTT, followed by dialysis into refolding buffer (10 mM Tris-HCl pH 7.5, 2 M NaCl, 1 mM EDTA and 5 mM β-mercaptoethanol). Refolded sample was subsequently purified by a Superdex 200 10/300 GL column to isolate the histone octamer.

### Nucleosome assembly for cryo-EM studies

For the nucleosomal DNA, we used a 158 bp DNA fragment amplified by PCR from a 200 bp G-block containing the human IGFBP1 promoter sequence (from -225 to -26) ordered from Integrated DNA Technologies (IDT). The DNA sequences for the G-block, amplified fragment and primers are shown below.

*IGFBP1 G-block DNA sequence (200 bp) (5’ → 3’):*

TTTAACTGAGGGCCTGAACCCCCTAACAACGGGACAAACAGTATGAAATCAAAACCGTTTACCCTC CTCCCACCAGCGGTTTGCGTAGGGCCTTGGGTGCACTAGCAAAACAAACTTATTTTGAACACTCAGC TCCTAGCGTGCGGCGCTGCCAATCATTAACCTCCTGGTGCAAGTGGCGCGGCCTGTGCCCTTTATAA

*IGFBP1 DNA sequence (158 bp, PCR amplified) (5’ → 3’):*

GGGCCTGAACCCCCTAACAACGGGACAAACAGTATGAAATCAAAACCGTTTACCCTCCTCCCACCA GCGGTTTGCGTAGGGCCTTGGGTGCACTAGCAAAACAAACTTATTTTGAACACTCAGCTCCTAGCGT GCGGCGCTGCCAATCATTAACCTCC

Primers Sequence (5’→3’):

Forward: GGCCTGAACCCCCTAA

Reverse: GGAGGTTAATGATTGGCAG

The amplified 158 bp IGFBP1 DNA sequence contains the cognate binding sites for FoxO1: 5’-TTATTTTGAA-3’, and for HNF1α: 5’-GCCAATCATTAAC-3’ transcription factors. Of note, a partial pseudo-binding site for exits at near the 5’-end of the sequence: 5’-**TGA**A**CC**C**CC**TAAC-3’ (the non-canonical bases are indicated in bold).

The histone octamer and nucleosomal DNA fragment were mixed in a 1:0.9 molar ratio in a buffer containing 20 mM Tris-HCl (pH 8) and 2 M NaCl. The mixture was placed into Slide-A-Lyzer MINI Dialysis Devices with a cut-off of 3.5 kDa (Thermo Scientific). The dialysis tube was then immersed into 1 L of buffer containing 2M NaCl, 20 mM Tris-HCl (pH 8) and 5 mM β-mercaptoethanol, and subjected to dialysis with buffer exchange against 2 L buffer containing 250 mM NaCl, 20 mM Tris-HCl pH 8 and 5 mM β-mercaptoethanol for 18h at 4 °C. The next day, the dialysis tube was immersed into the remaining volume of the low-salt buffer to continue dialysis for 5h at 4 °C. After that, the dialysis tube was immersed into 2 L of buffer containing 20 mM NaCl and 20 mM HEPES (pH 8) to dialyze overnight at 4 °C. Finally, the nucleosomes were analyzed by 5% native PAGE using SYBR Gold staining (Figure S1a).

### FoxO1 and HNF1a expression and purification

FoxO1 (residues 144-266) and HNF1α (residues 90-288) purification was performed by cloning the gene encoding the corresponding construct, into a vector containing a 6×histidine-SUMO tag and the recombinant proteins were expressed in *Escherichia coli* Rosetta cultured at 18 °C and induced with 0.2 mM IPTG overnight. Cells were harvested and lysed by sonication on ice in lysis buffer containing 500 mM NaCl, 20 mM HEPES (pH 7.5), 10 mM imidazole, 10% (v/v) glycerol, and protease inhibitor cocktail. Cell debris was removed by centrifugation, and the clarified supernatant was incubated with Ni–NTA agarose beads (Cytiva) for 3h at 4 °C. The beads were washed with 30 mL of lysis buffer, and bound proteins were eluted with elution buffer containing 150 mM NaCl, 20 mM HEPES (pH 7.5), and 500 mM imidazole. The SUMO tag was removed by ULP1 protease digestion, and the sample was further purified by Heparin affinity chromatography (Cytiva) using a linear NaCl gradient from 150 mM to 2 M in 20 mM HEPES (pH 7.5) and 10% glycerol. Target fractions were pooled, concentrated to 0.5 mL, and subjected to size-exclusion chromatography on a Superdex 75 Increase 10/300 GL column. Following size-exclusion chromatography (Figure S2), proteins were concentrated to ∼0.5 mg/mL, flash-frozen in liquid nitrogen, and stored at -80 °C.

### Binding assays

Binding assays for both FoxO1 and HNF1α were performed using electrophoretic mobility shift assays (EMSA) on a 5% native PAGE. The nucleosome concentration ([Nuc]) was maintained at 300 nM and the FoxO1 or HNF1α concentrations varied according to the molar ratios tested. The gel was then assessed using SYBR Gold staining (Figure 1e and Figure 2b in main text).

### Cryo-EM sample preparation

Samples of nucleosomes in complex with FoxO1, HNF1α, or both factors, were assembled using 1.5 µM of *IGFBP1* nucleosome and 10× molar excess of FoxO1 or HNF1α. Samples were preincubated at room temperature for 10 minutes in buffer containing 20 mM HEPES pH 8 and 20 mM NaCl. Then, 3.5 µl of the sample were deposited on Quantifoil 1.2/1.3 holey carbon gold grids (previously glow-discharged using a Tergeo-EM plasma cleaner, PIE Scientific), blotted for 10 seconds with a blot force of 10, and plunge-frozen in liquid ethane using a Vitrobot Mark IV (Thermo Fisher Scientific, USA) at 22°C and 100% humidity.

### Cryo-EM data collection

Cryo-EM Dataset 1 and Dataset 2 were acquired using a 200 kV Talos Arctica cryo-electron microscope (Thermo Fisher Scientific) at 36,000× magnification, collected in super-resolution counting mode (super-resolution pixel size of 0.5705 Å/pixel) on a K3 direct electron detector (Gatan). Cryo-EM Dataset 3 (non-tilted and tilted +/-30°) was acquired using a 300 kV Titan Krios G3 cryo-electron microscope (Thermo Fisher Scientific) using a K3 direct electron detector (Gatan) at 81,000× magnification in super-resolution counting mode (super-resolution pixel size 0.524 Å/pixel). All data sets were collected with a total exposure dose of 50 *e*^−^/Å^2^ fractionated across 50 frames during movie frame recording, and defocus values ranging from −0.8 µm to −1.8 µm. All data sets were automatically collected using SerialEM^2^.

### Cryo-EM image analysis

Data processing of all the cryo-EM datasets was performed using cryoSPARC v4.7.1^3,4^ and RELION5 software^5,6^. For simplicity, we will describe in detail the data analysis workflow followed for Dataset 1, which resulted in the elucidation of the Free ^IGFBP1^Nuc Class I, Free ^IGFBP1^Nuc Class II, and ^IGFBP1^Nuc–FoxO1 structures (Figure S3). We note that the analysis for Dataset 2 and Dataset 3, resulting in ^IGFBP1^Nuc–HNF1α and ^IGFBP1^Nuc–FoxO1–HNF1α structures, respectively, was performed following similar workflows and we detail the differences as well (Figure S4 and S5).

### Dataset 1 processing

We assembled the Nucleosome-FoxO1 complex and prepared cryo-EM grids as described above. From this sample, we collected Dataset 1 consisting of 12,979 movie frames, which were then aligned using Patch Motion Correction within cryoSPARC. Defocus estimation and contrast transfer function (CTF) fitting were performed using Patch CTF Estimation. In the corrected micrographs, we could readily observe particles with the size and features expected for nucleosomes (Figure S3). Particle picking was then performed using the Blob Picker algorithm, followed by multiple rounds of 2D classification/particle selection to obtain a clean subset of particles. Then, 400,000 particles were randomly selected to perform an ab initio 3D reconstruction (N=3), from which one volume was selected to perform Heterogeneous Refinement (N=4). In the obtained 3D classes, we can clearly distinguish the co-existence of free nucleosome and distorted nucleosome populations. From this point, we split the data processing in two pathways: the orange branch, focused in analyzing the free nucleosomes’ population, and the green branch, which will analyze the distorted nucleosome classes. From the orange branch, and Homogeneous Refinement followed by a Heterogeneous Refinement (N=3) round, allowed us to separate individual nucleosome structures from the stacked nucleosomes populations, which were also visible in our individual micrographs (Figure S6). A final round of 3D classification without alignment and Non-uniform Refinement, allowed us to obtain two distinct Free IGFBP1 nucleosome structures: Class I, fully wrapped at 3.5 Å overall resolution, and Class II, partially unwrapped at 3.7 Å overall resolution (Figure S3 and S6). From the green branch, an Heterogeneous Refinement round (N=4) allowed us to separate different populations corresponding to stacked nucleosomes and individual distorted nucleosomes, where these last ones were refined separately. Later, a 3D classification with alignment process resulted in different distorted nucleosome classes, where one class showed an extra attached small density after being subjected to homogenous refinement. We then employed a mask surrounding the extra density and performed focused 3D classification without alignment (N=10). One of the classes was selected and refined to 6.2 Å overall resolution and showed a clear distorted nucleosome, harboring an extra density attached to it attributed to bound FoxO1 (Figure S3 and S6).

### Dataset 2 processing

We assembled the Nucleosome-HNF1α complex and prepared cryo-EM grids as described above. From this sample, we recorded Dataset 2 consisting of 10,469 movie frames in total (collected in two parts), which were then aligned using Patch Motion Correction within cryoSPARC. Defocus estimation and contrast transfer (CTF) fitting were performed using Patch CTF Estimation in cryoSPARC. Subsequent steps were performed similarly to Dataset1 to clean the particle subset and generate the initial reconstructions. Of note, representative 2D class averages in this data set readily reveal an extra density bound to the nucleosome specifically in top-view orientations. An intermediate 3D map obtained by Homogeneous refinement from ∼318K particles, was resolved at 3.5Å overall resolution and shows an ill-defined density bound to the nucleosome. These particles were then exported to Relion software to perform 3D refinement. The resulting 3D map shows a well-defined nucleosome at high-thresholds, while the attached small density (named here as lobe A), appears fuzzy at lower-thresholds. We defined a mask around this region and followed a pipeline with a combination of Particle subtraction, particle re-boxing, and 3D classification without alignment (N=3, T=500). We then isolated a stable class (containing∼94k particles) where the lobe A density reveals more features. Then, this particle population underwent revert subtraction to perform a final 3D refinement resulting in a 4Å overall resolution map where the lobe A appears clearer. A similar class of a nucleosome with an extra-density bound to it was identified in a sub-population (∼162K particles) from Dataset 3 (see Class B, Figure S4), then these two classes were combined. The combined ∼256K particles were subjected to a 3D refinement resulting in a 3.1Å overall resolution map. At high-threshold, this 3D reconstruction reveals a well-resolved nucleosome clearly fully wrapped in one DNA arm, while the other DNA arm appears unwrapped and harbors the fuzzy lobe A bound to it, visible at lower-thresholds. We designed a mask involving the stable nucleosome region and performed Particle Subtraction followed by 3D refined, resulting in a 3D reconstruction Nucleosome at 3Å overall resolution, where the IGFBP1 DNA sequence was assigned unambiguously, and we determined that the proximal DNA arm remains wrapped while the distal DNA appears unwrapped. In parallel, we designed a mask encompassing the lobe A and including its DNA-interacting region, and performed Particle Subtraction, including re-center in mask and re-boxing in a smaller 180x180 pixel box size. We then back-projected the particles to get a new 3D reference to perform 3D classification without alignment (N=4, T=500). A single population harboring ∼49K particles, exhibited clear secondary structural features, and was then subjected to 3D refinement resulting in a final 3D reconstruction solved at 4.3Å overall resolution. At this resolution, we clearly distinguish that an HNF1α dimer interacts with the distal DNA arm region, causing significant nucleosome unwrapping as observed in the composited cryo-EM map (Figure S4 and Figure S6).

### Model building, refinement and validation

The final refined cryo-EM maps obtained for all the structures described above, were then B-factor and DeepEMhancer^7^ sharpened before the initial atomic coordinate modeling.

For ^IGFBP1^Nucleosome Class I and Class II, the initial coordinates models were obtained using *ModelAngelo*^8^ (within RELION5) to indicate the corresponding protein and nucleic acids sequences. For the ^IGFBP1^Nucleosome-Fox structure, the initial model were obtained by rigid-body fitting of the ^IGFBP1^Nucleosome Class I atomic coordinates into the corresponding density map using UCSF ChimeraX^9^. For the ^IGFBP1^Nucleosome-HNF1α complex, the starting coordinate model was obtained in two steps. First, , the initial coordinates for the nucleosome-focused region were obtained by rigid-body fitting of the ^IGFBP1^Nucleosome Class I atomic model. Second, the initial model for the focused HNF1α-DNA region was obtained by rigid-body fitting the atomic model of the human HNF1α dimer bound to its cognate linear DNA target (PDB 1IC8)^10^. All initial atomic models were then iteratively rebuilt in COOT^11^ and refined using the real space refinement program in PHENIX^12^.

All validation and refinement statistics are shown in Table S1.

### Nucleosome assembly and purification for Optical Tweezers experiments

Recombinant *Xenopus laevis* Histone octamers and the 158 bp IGFBP1 sequence, flanked by 20 bp and 10 bp DNA handles on each side, were combined at a ratio of 1:1 in 500 µL of high-salt buffer (10 mM Tris-HCl pH 8.0, 2 M NaCl, 1 mM EDTA, 0.5 mM DTT, and 1 mM PMSF) at a final concentration of 100 ng/µL of DNA. Assembly solutions were dialyzed against 500 mL of high-salt buffer for 1h at 4 °C, followed by a linear gradient dialysis against 2 L of low-salt buffer (10 mM Tris-HCl pH 8.0, 1 mM EDTA, 0.5 mM DTT, 1 mM PMSF) using a peristaltic pump with a 0.8 mL/min flow rate and continuous stirring. A final dialysis of 3 h in 500 mL of low salt buffer was done to reduce the residual NaCl concentration. The nucleosome reconstitution was checked by 5% acrylamide native electrophoresis using 0.2X TBE (Tris-borate-EDTA) as a running buffer. and fractions containing nucleosomes were concentrated by centrifugation using 100K Amicon Ultra filters (Millipore). After concentration, samples were dialyzed against HE buffer (20 mM HEPES-NaOH pH 7.5, 1 mM EDTA, 1 mM DTT) and stored at 4°C.

### Oligo-bead preparation

We conjugated double-stranded oligonucleotides (IDT) to 1 um carboxy functionalized polystyrene beads (Bangs lab) using the protocol found in Gabizon et al^13^. We took 53 µl of 10% w/v of 1 µm carboxyl Polystyrene beads and centrifuged at 5000 g at 4°C for 8 min. Beads were washed with 1 ml of ultrapure water and 0.75 ml of 200 mM MES buffer and resuspended in 106 µl of MES. The crosslinking reaction was carried out with 100 µl of washed beads with 50 µl of dsOligo, and 30 µl of 2M of 1-ethyl-3-[3-dimethylaminopropyl]carbodiimide (EDC). Reaction was incubated at 20°C overnight at 1200 rpm. Next day, we stopped the reaction with 460 µl of 1M Tris pH. 8.0. Beads were washed with 1 ml of 200 mM MES and 1 ml of TE8 buffer (10 mM Tris pH 8.0 + 1 mM EDTA). We added 200 µl of 10 mg/ml of Casein (Sigma-Aldrich) and incubated at 22°C for 15 min and 10-20 sec of vortex every 2 min. We washed the casein with 1 mL of TE8 buffer. Finally, beads were resuspended in 500 µl of TE8/5 buffer (20 mM Tris pH 8.0 + 5 mM EDTA + 10mM sodium azide) and 116 µl of 80% glycerol. Aliquots were frozen with liquid nitrogen and stored at -80°C. Aliquots ready to use for optical tweezers experiments were kept at 4°C.

### Sample preparation for optical tweezers experiments

Purified single nucleosomes were ligated to 570 bp (5’-end) and 700 bp (3’-end) DNA handles at a concentration of 50 nM each, using E. coli DNA ligase (NEB) for 2h at 16 °C in the presence of 0.02% NP-40. Later, we attached nucleosomes to the Oligo-beads using E. coli DNA ligase at 16°C for 2 h via the 4 nt sticky end of the 570 bp handle of the ligation product above. The other 700 bp handle was designed with a 5’-biotin modification to allow for later attachment to streptavidin-coated polystyrene bead (Banglabs).

### Single-molecule optical tweezers set-up

We used a homebuilt dual-trap high-resolution optical tweezers instrument^14^. Data was collected at 2.5 kHz, with a trap stiffness of ∼0.35 pN/nm, at room temperature and in the pulling buffer (20 mM NaCl and 20 mM HEPES-KOH pH 8.0). To avoid photodamage and prevent contamination, the buffer was supplemented with 10 mM sodium azide, and we used the Steriflip® system for filtering and degassing the buffer for 1 hour before performing experiments^15^. All experiments were carried out in a homebuilt glass microfluidic chamber with three channels equilibrated with pulling buffer. We performed all pulling/relaxing measurements in the middle channel. Also, before introducing the 1 um polystyrene beads with nucleosomes, the top channel was passivated for 30 min with 20 mM HEPES-KOH pH 7.5, 50 mM KOAc, 1 mg/mL BSA (NEB), followed by a second passivation for 30 min with 20 mM HEPES-KOH pH 7.5, 50 mM KOAc, 0.5% pluronic acid^15^. We introduced the passivated 1 um streptavidin beads in the bottom channel. Each nucleosome and streptavidin bead were dispensed via a microchannel (dispenser) connecting the top and bottom channel with the middle one. Each bead was captured in each optical trap and moved away 1 mm from both dispensers to prevent flow artifacts and reduce noise.

### Single-molecule optical tweezers data collection

We controlled the instrument using a custom software written in LabVIEW^16^. Each pair of beads was calibrated to estimate the off-set and trap stiffness via the power spectrum method and fitting to the Lorentzian equation^15,17,18^. Tethering was performed by moving the 1 µm streptavidin-passivated bead close to the 1 um nucleosome-ligated bead. Later, we slowly increased the trap separation by moving the streptavidin bead away. A tether between beads is recognized by the slow, monotonic increase of force below 2 pN. Because the total length of the molecule between both beads must be ∼440 nm, we only considered tethers of 440 ± 20 nm to perform further measurements. Pulling/relaxing cycles per molecule were performed at 20 nm/s in pulling buffer. Per molecule, we recorded extension, tether length and time. Single tethers were identified when their rupture occurred in a single step with the force dropping to 0 pN, or by observing the dsDNA overstretching force (∼65 pN). Only single tethers were selected for analysis.

### Analysis of optical tweezers data

Each molecule was analyzed individually using a custom MATLAB code^15,19^. We measured the force at which a transition occurred as well as the corresponding change in extension for the first and second pulling. We only selected molecules whose first pulling was an octasome, which is identified by the low-force transition and high-force transition. Other sub-nucleosomal particles are identified by a cooperative transition above 15 pN (hexasome) with a change in extension similar to that observed in the high-force transition of the octasome^15^. Tetrasomes are identified by having a non-cooperative transition at forces below 5 pN^15^. The probability of disassembly of octasomes to other sub-nucleosomal particles (hexasomes/tetrasomes) or to bare DNA was calculated as the fraction of observing a specific disassembly event (e.g. octasome in first pulling to hexasome/tetrasome in second pulling) divided by the total number of first to second pulling cycles. Although all molecules were pulled/relaxed many times till we observe full conversion to bare DNA, only the first and second pulls were analyzed to calculate disassembly probabilities. Sometimes, the tether broke after observing the high-force transition of the octasome in the first pulling. These molecules were included in the force transition analysis but excluded from the disassembly probability analysis. Tethers that ruptured after the second pulling were included in the disassembly probability analysis. Because the low-force transition of the octasome was difficult to measure, we aligned all the first pulling traces of octasomes per condition at 8 pN and determined their average behavior.

### IDLI data analysis

IDLI analysis was carried out as in Abdulhay et al^20^. Nucleosome footprints were sampled from d6 endoderm SAMOSA data (also from^20^), such that footprints fell within ENCODE-defined DHS from human hepatocytes and contained FoxO1 and/or HNF1α binding motifs as defined by Vierstra *et al*^21^. Nucleosome types were further classified into groups using the hclust package in R, and horizon plots, nucleosome footprint sizes, and intranucleosomal accessibility patterns were visualized using scripts available at https://github.com/RamaniLab/IDLI. Following nucleosome type definition, custom Python scripts were used to compute relative enrichment and depletion of nucleosome types for nucleosome footprints containing FoxO1 motifs only, HNF1α motifs only, or the combination of both motifs (maximum distance between motifs = 50 nucleotides). All Fisher’s exact *p* values were then corrected to account for multiple hypothesis tests using Storey’s *q* value^22^.

## ACKNOWLEDGMENTS

We thank Kedar Sharma, D. Toso and R. Thakkar at the Cal-Cryo EM facility at QB3-Berkeley for help with EM data acquisition, and to P. Tobias and K. Stine for computing support. We also thank the valuable discussions and feedback from members of the Bustamante and the Nogales labs. Z.Y. acknowledges that he is presently an investigator of the SUSTech Institute for Biological Electron Microscopy.

## AUTHOR CONTRIBUTIONS

A.J.F.A., C.B. and E.N. conceptualized and designed the study. A.J.F.A. and Z.Y. performed protein production and nucleosome assemblies for cryo-EM and biochemical assays. AJ.F.A. performed cryo-EM sample preparation, cryo-EM data collection, data analysis, atomic modeling and structural interpretation. C.D.C. performed nucleosomes assembly and DNA handle constructs designs to carry out optical tweezer’s experiments. R.P.S. performed optical tweezers data collection, data analysis and interpretation. H.J.R. and V.R. produced the human endoderm differentiated cells and performed footprinting data collection and analysis. A.J.F.A. wrote the original draft of the manuscript with the contribution, revision, editing, and approval of all the authors.

## DATA AND CODE AVAILABILITY

The atomic coordinates and cryo-EM maps for the IGFBP1 nucleosome Class I and II, IGFBP1 nucleosome-FoxO1 and IGFBP1 nucleosome-HNF1a structures have been deposited in the PDB and EMDB with identifiers PDB 39XH, PDB 39XX, PDB 39XY, PDB 39YB, EMD-79868, EMD-79878, EMD-79879, and EMD-79882, respectively.

## SUPPLEMENTAL FIGURES

**Figure S1:**
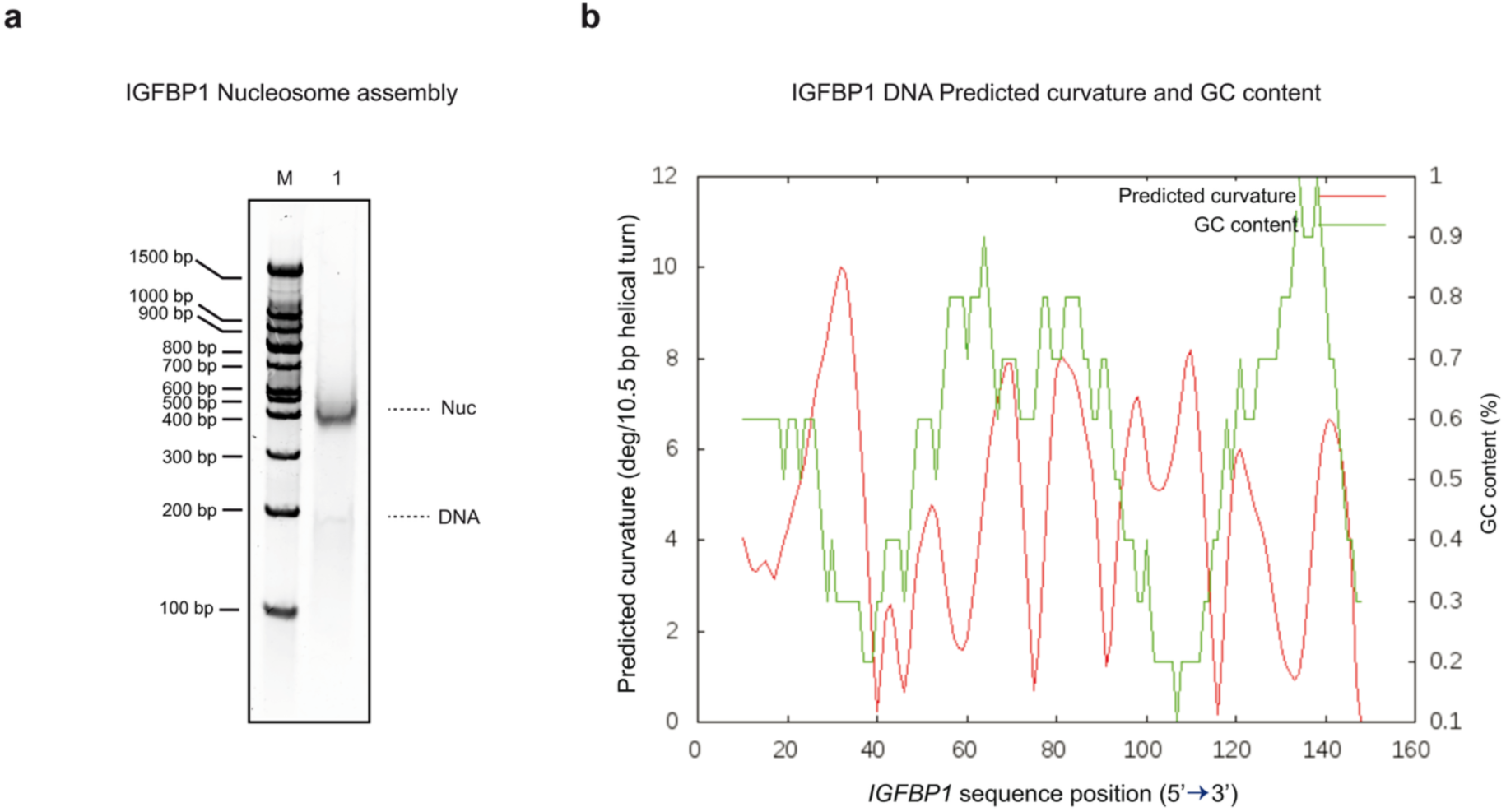
Assembly of the IGFBP1 nucleosomes and IGFBP1 curvature DNA analysis, related to Figure 1. **a.** Native Tris base, boric acid, and EDTA (TBE) gel, TBE 5%, showing the optimized assembly of the *IGFBP1* nucleosome using an octamer:DNA molar ration of 1:0.9. **b.** Analysis of the Predicted curvature (deg/10.5 bp helical turn) and GC content (%) along the IGFBP1 DNA sequence. The calculations and plot were performed using the bend.it server^23^.

**Figure S2:**
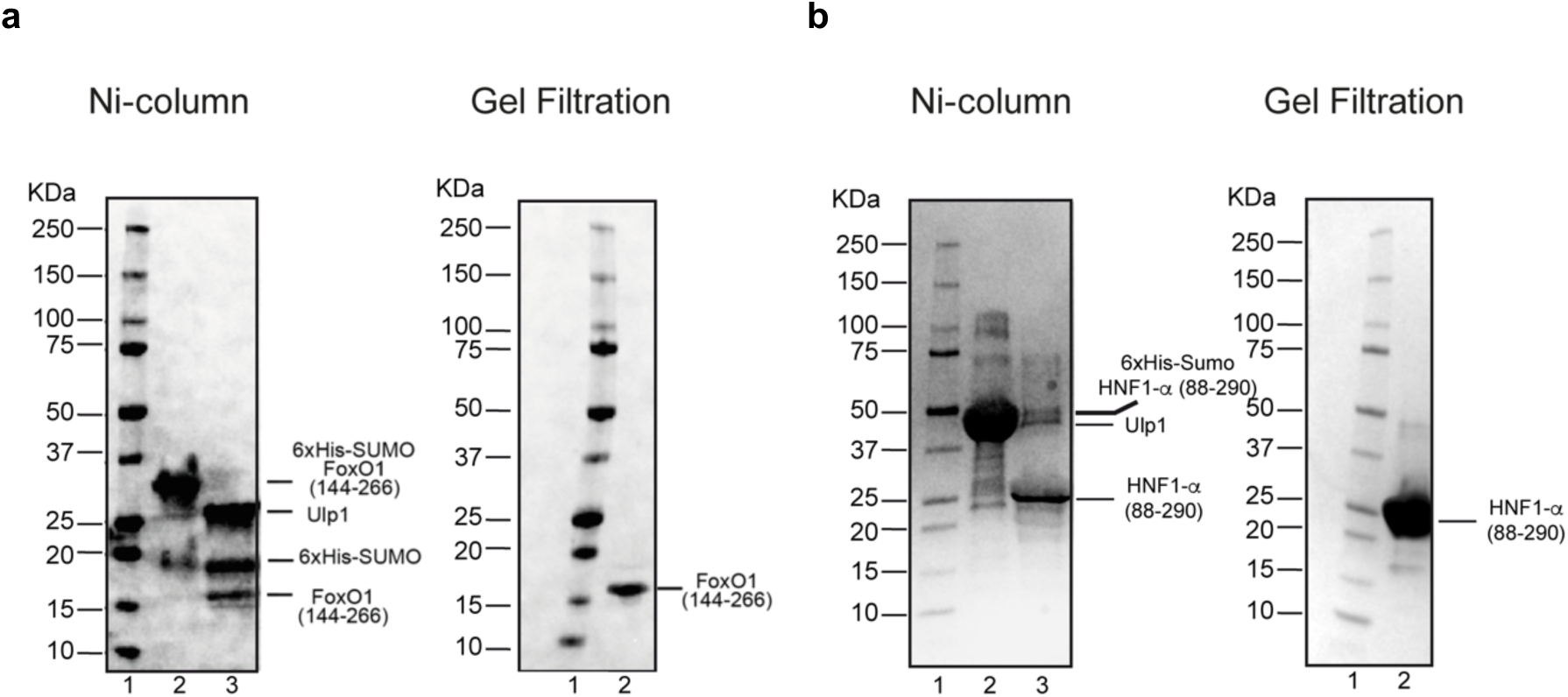
Protein purification process for FoxO1 and HNF1α, related to Figure 1 and 2. **a.** (Left) SDS-PAGE analysis of the human FoxO1 (residues 144-266) protein containing 6xHis-SUMO tag after purification by Ni-column. Lane 1: marker, lane 2: 6xHis-SUMO-FoxO1 fusion protein before tag cleavage, lane 3: FoxO1 (residues 144-266) after 6xHis-SUMO tag cleavage using Ulp1. (Right) SDS-PAGE analysis after purification of FoxO1 (residues 144-266) by size-exclusion chromatography (Gel Filtration). Lane 1: marker, lane 2: FoxO1 (residues 144-266). **b.** (Left) SDS-PAGE analysis of the human HNF1α (residues 88-290) protein containing 6xHis-SUMO tag after purification by Ni-column. Lane 1: marker, lane 2: 6xHis-SUMO-HNF1α fusion protein before tag cleavage, lane 3: HNF1α (residues 88-290) after 6xHis-SUMO tag cleavage using Ulp1. (Right) SDS-PAGE analysis after purification of HNF1α (residues 88-290) by size-exclusion chromatography (Gel Filtration). Lane 1: marker, lane 2: HNF1α monomer (residues 88-290).

**Figure S3:**
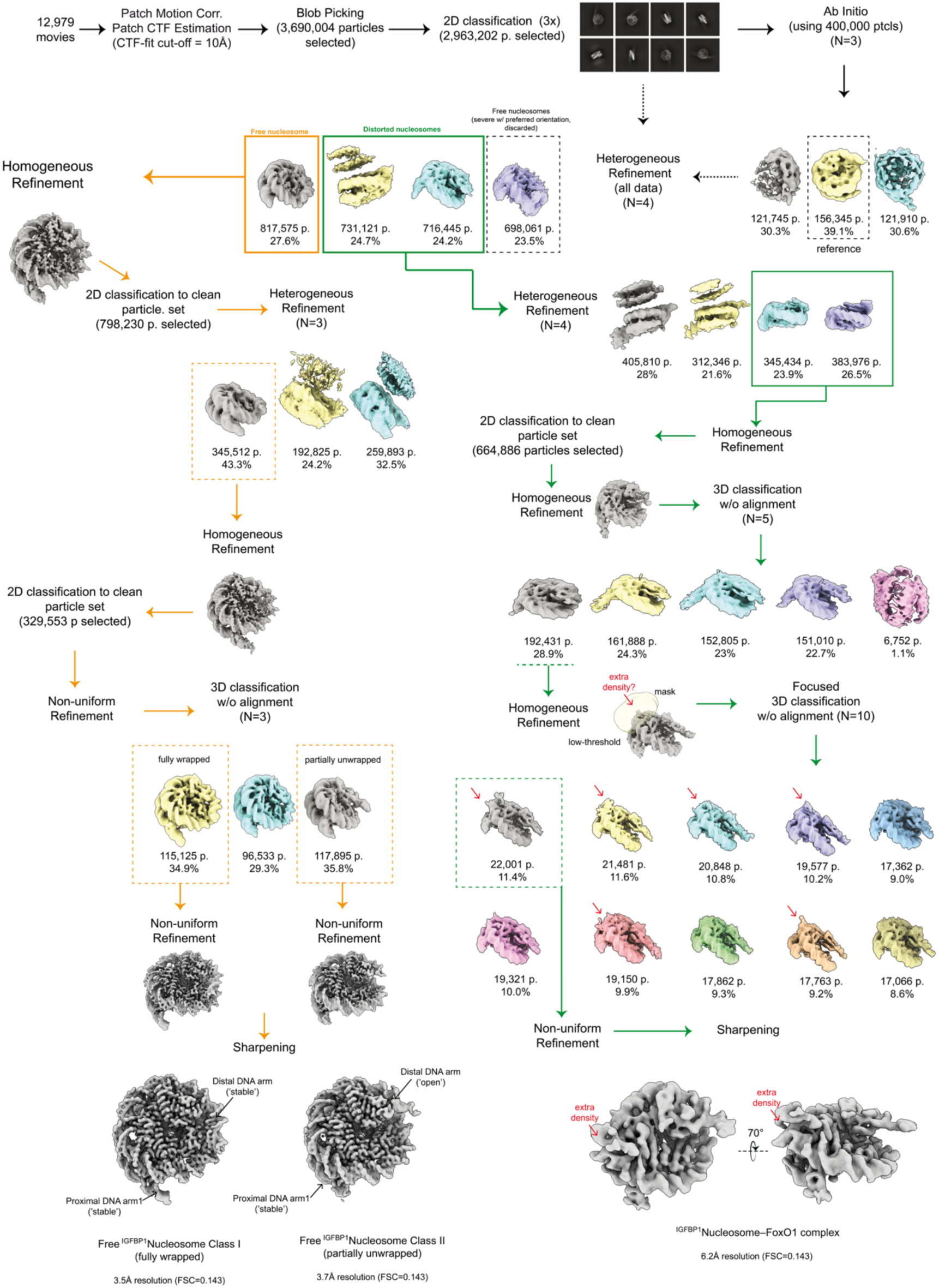
Cryo-EM processing workflow used for data set 1 to obtain the structures of the IGFBP1 Nucleosome, alone (^IGFBP1^Nuc Class I and Class II) or bound to FoxO1 (^IGFBP1^Nuc–FoxO1), related to. Figure 1.

**Figure S4:**
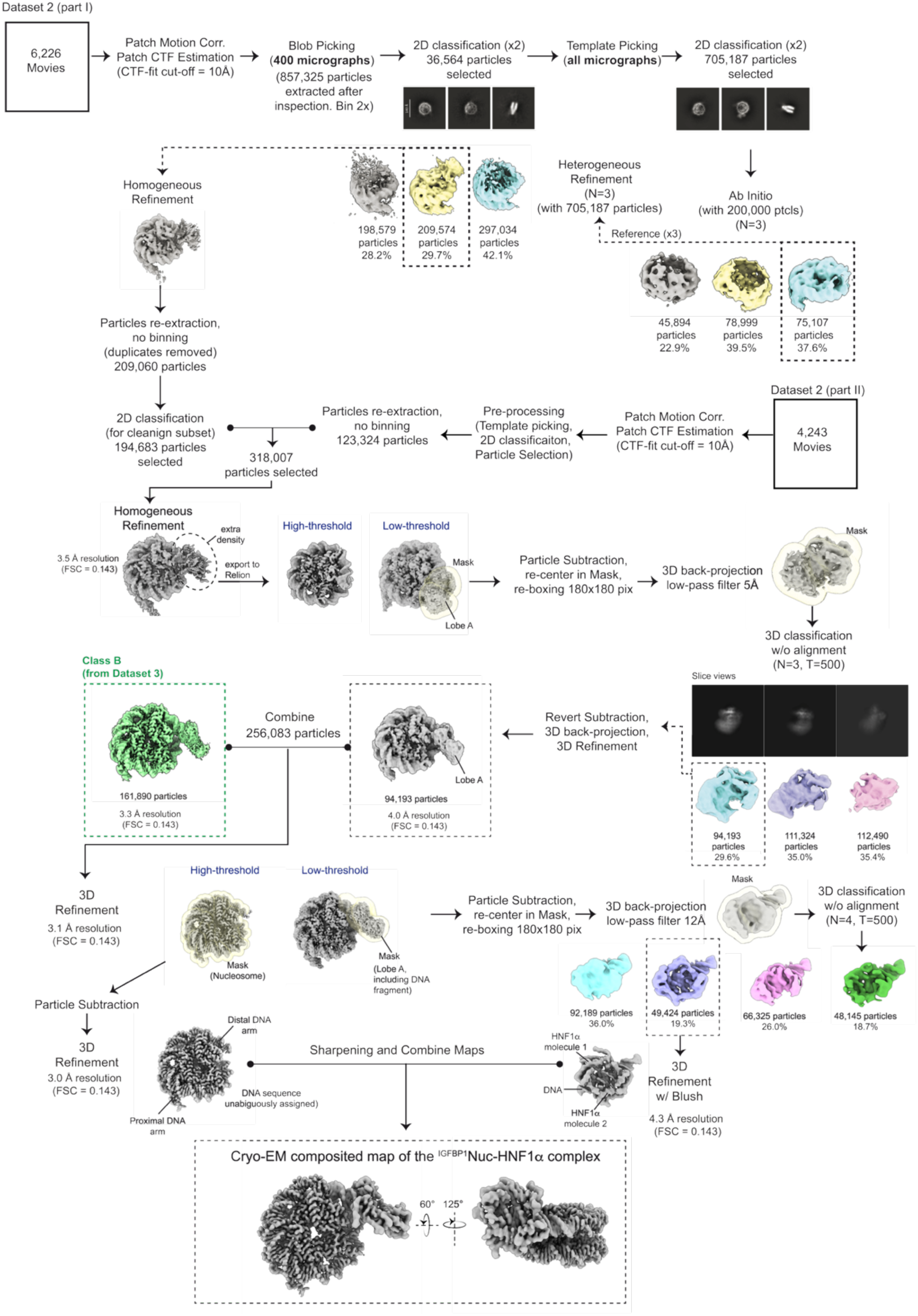
Cryo-EM processing workflow used for data set 2 to obtain the structure of the *IGFBP1* Nucleosome bound to HNF1α (^IGFBP1^Nuc–HNF1α), related to Figure 2.

**Figure S5:**
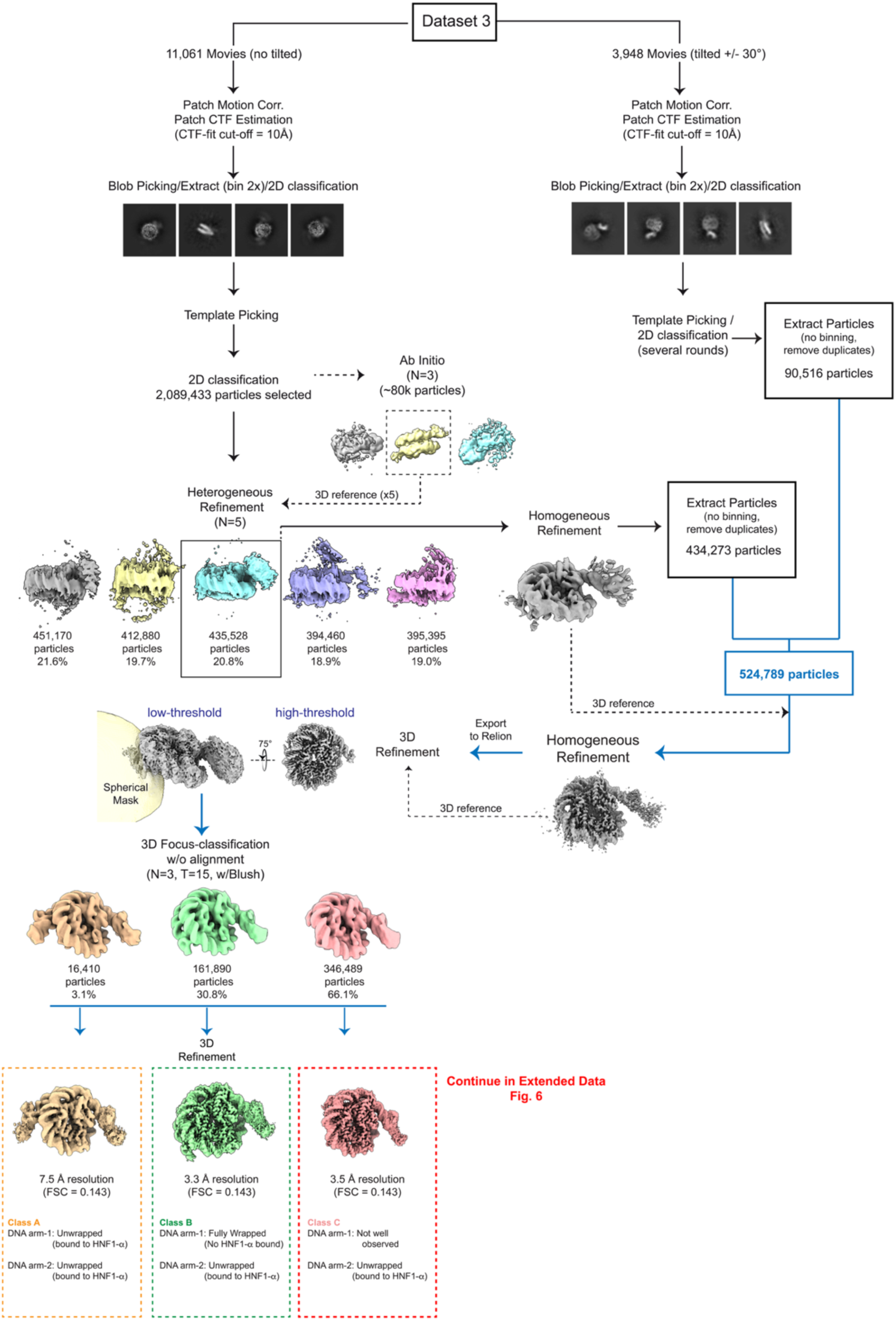
Part I of the cryo-EM processing workflow used for data set 3 corresponding to the sample containing ^IGFBP1^nucleosome, FoxO1 and HNF1α, related to Figure 2.

**Figure S6.**
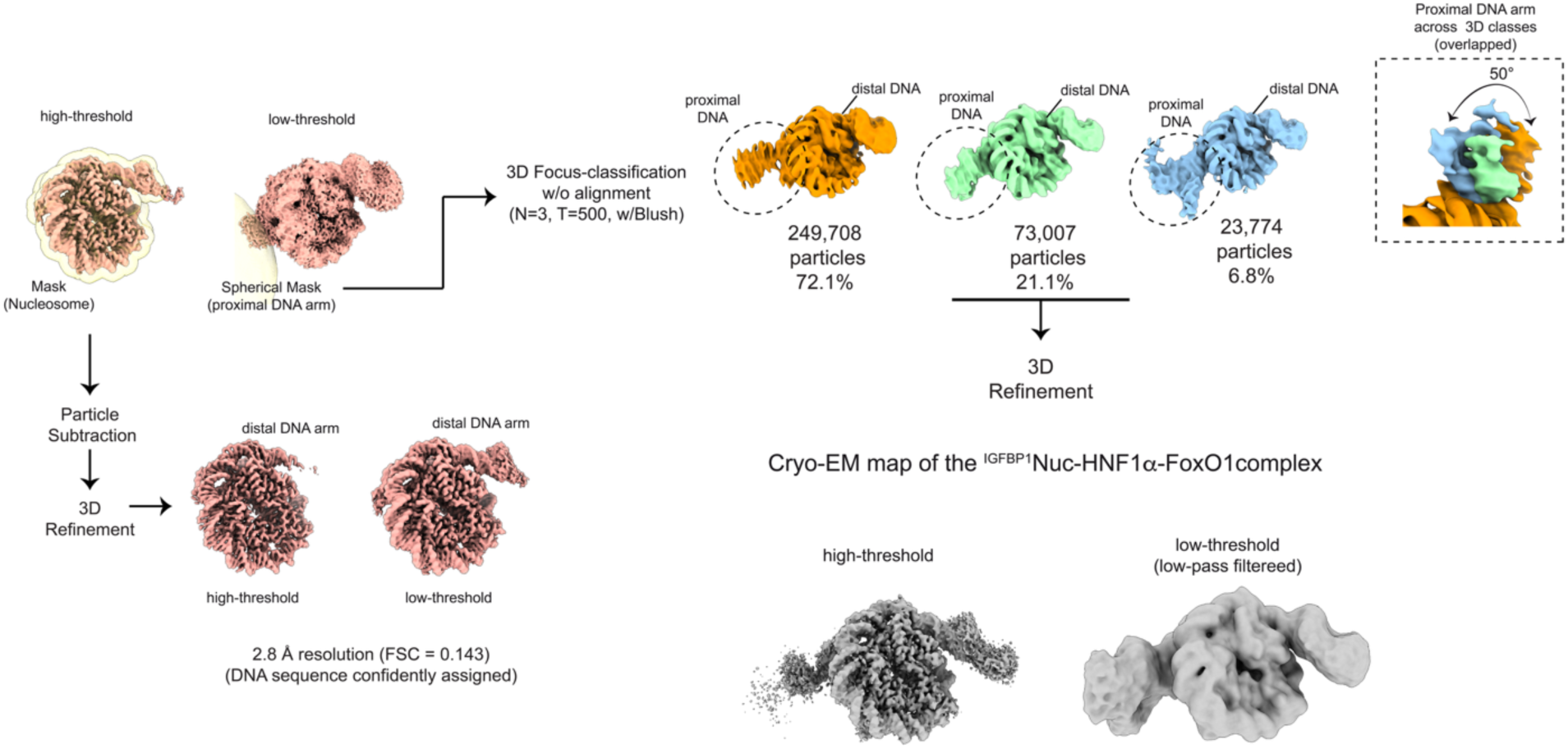
Part II of the cryo-EM processing workflow used for data set 3 corresponding to the sample containing ^IGFBP1^nucleosome, FoxO1 and HNF1α, related to Figure 2.

**Figure S7:**
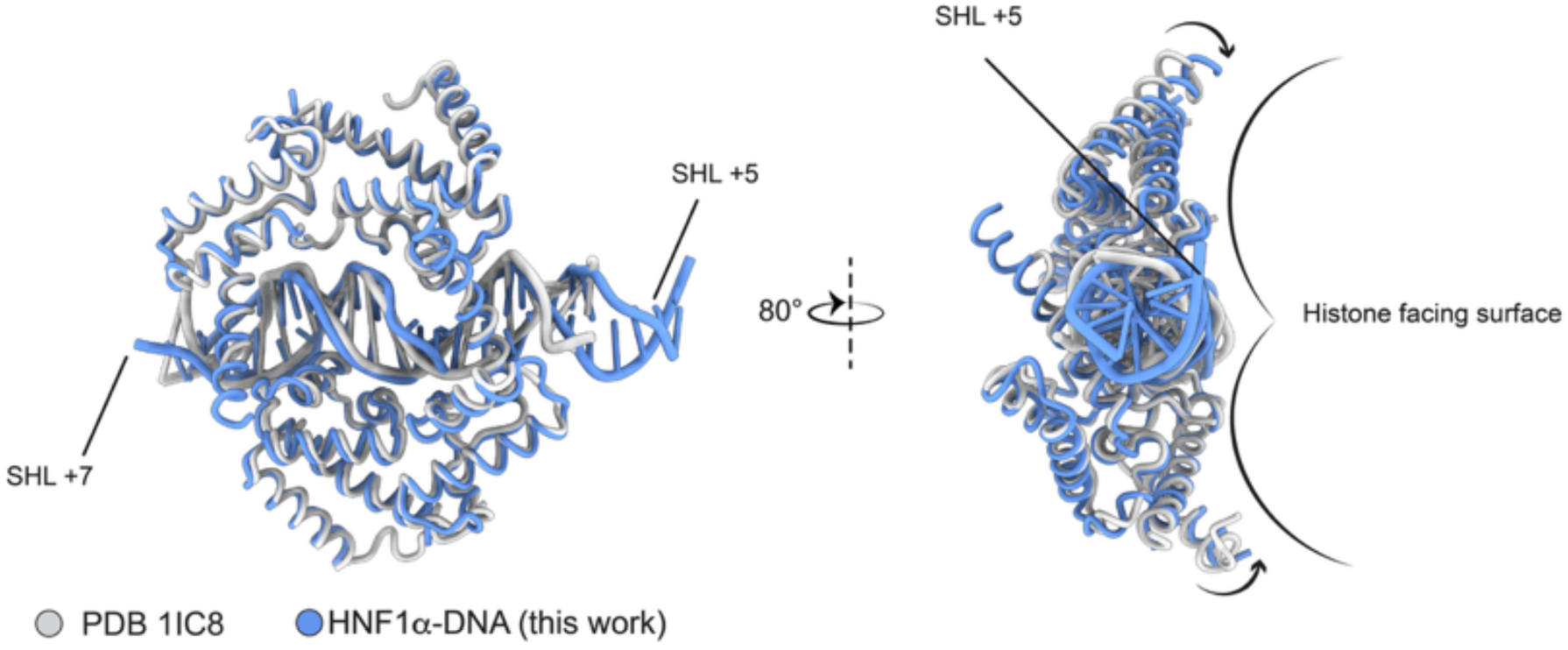
Comparison of the HNF1α dimer positioning in the ^IGFBP1^Nucleosome-HNF1α structure relative to its linear DNA-bound X-ray structure (PDB 1IC8), related to. Figure 2

**Figure S8:**
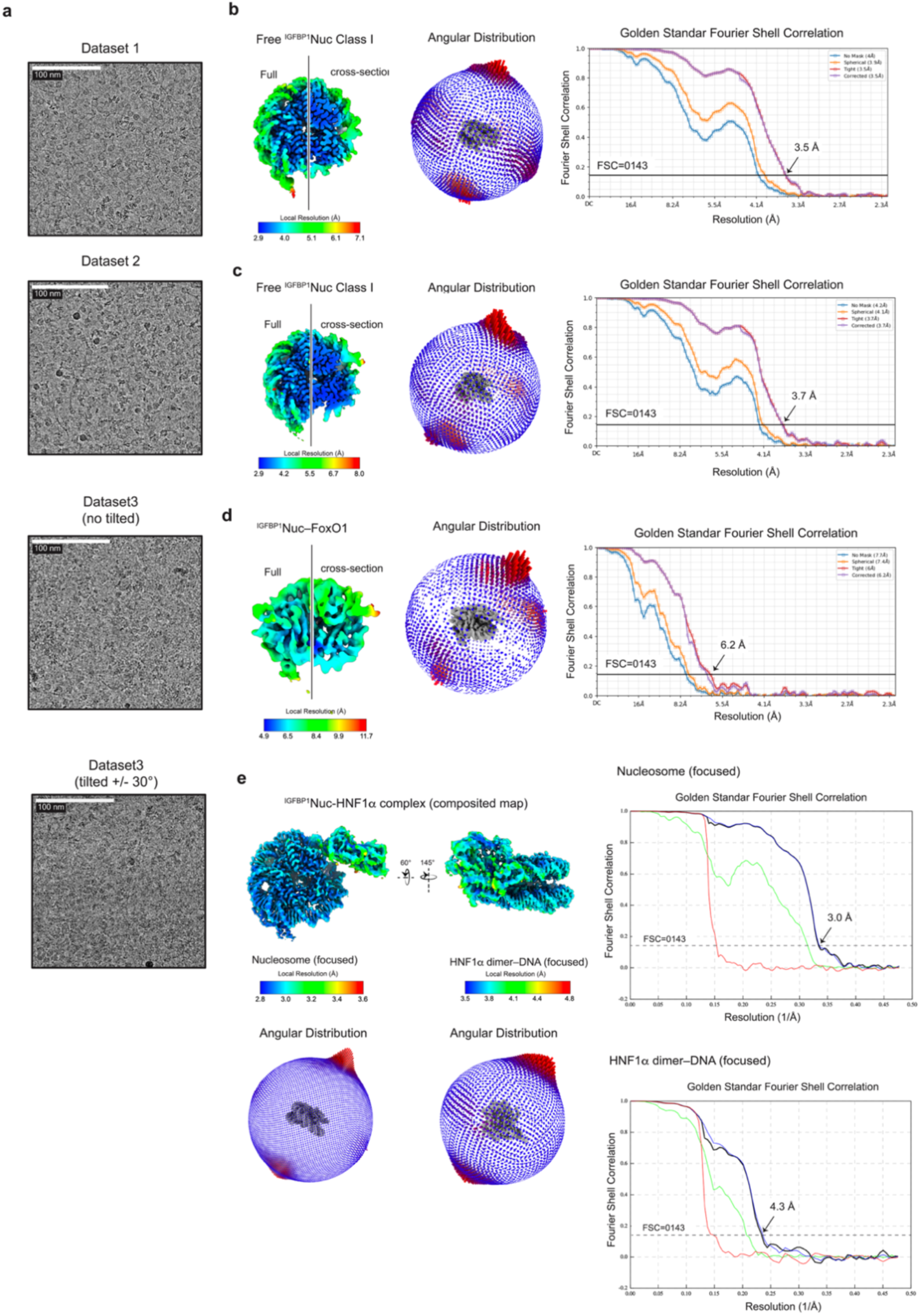
Cryo-EM data analysis statistics, related to Figure and 2. **a** Examples of cryo-EM corrected micrographs for the datasets analyzed. Scale bars are indicated. **b–f** Unsharpened cryo-EM map colored by local resolution as indicated in the color key (left), distribution plot (middle), and Fourier Shell Correlation (FSC) map vs map and map vs model plots (right) for the ^IGFBP1^Nuc Class I (b) and Clas II (c), ^IGFBP1^Nuc-Fox (d) and ^IGFBP1^Nuc-HNF1a (e) structures. For the local resolution plots, the line separates a cross-section on the left and a view of the cryo-EM map surface on the right.

**Figure S9:**
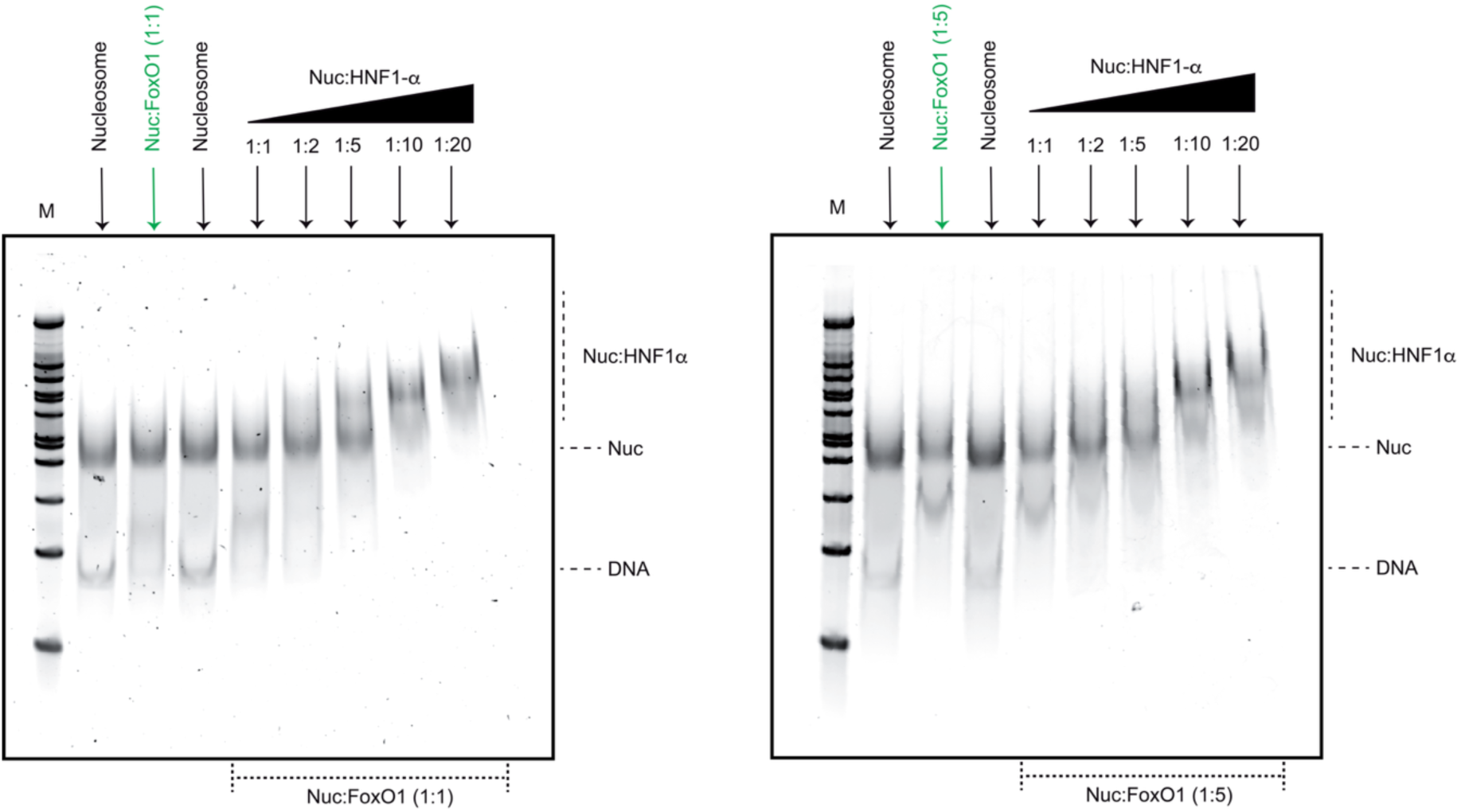
Cooperative binding analysis of FoxO1 and HNF1a to the IGFBP1 nucleosome, related to. Figure 2 EMSAs in which IGFBP1 nucleosomes were pre-incubated with FoxO1 at 1:1 (left) or 1:5 (right) molar ratios prior to the addition of HNF1a at different molar ration. Nucleosomes concentration was kept fixed at 300 nM.

**Figure S10:**
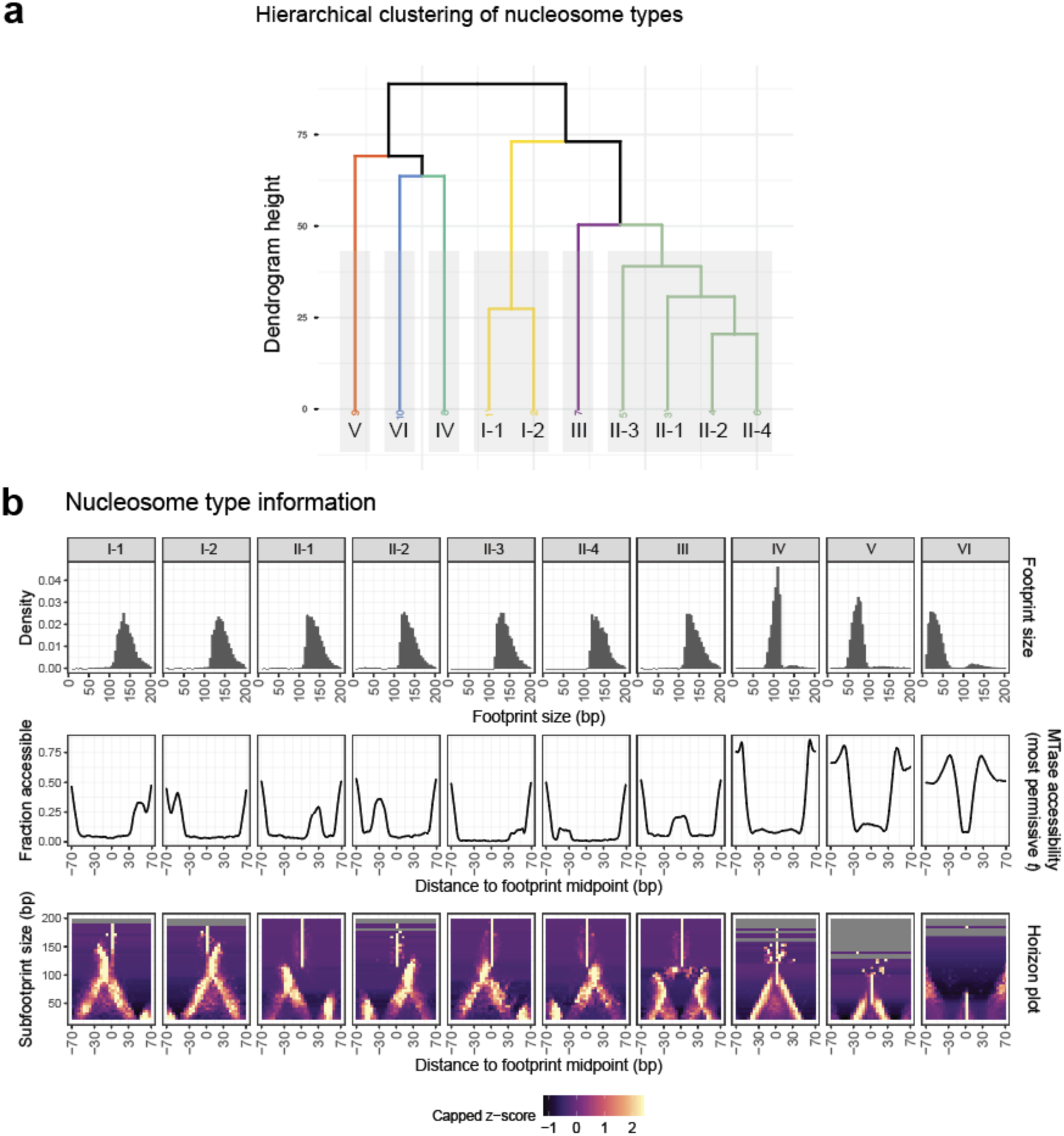
Technical details from IDLI analysis of FOXO1 or HNF1b motif containing nucleosome footprints in iPSC-derived endoderm, related to Figure 4. **a** Hierarchical clustering dendrogram demonstrating the grouping of specific nucleosome types into larger ‘groups’, as in^20^. **b** Top row: Footprint sizes for each nucleosome type called using the IDLI pipeline; Middle row; Average EcoGII methyltransferase accessibility at the most permissive transition probability t for each nucleosome type. This represents the highest resolution sequence-context corrected methyltransferase accessibility called for each nucleosome type. Bottom row: Horizon plot heatmap visualization for each nucleosome type, which visualizes the positioning and enrichment of successively caller ‘subfootprints’ that are called within each larger nucleosome footprint using the IDLI pipeline. Each horizon plot represents a structurally distinct nucleosome/subnucleosomal species defined using the neural network-hidden markov model (NN-HMM) framework employed by IDLI.

**Table 1:** Cryo-EM data collection, refinement, and validation statistics.

|  | IGFBP1 Nucleosome |  | IGFBP1 nucleosome–FoxO1 complex<br>(EMD-79879)<br>(PDB 39XY) | IGFBP1 nucleosome–HNF1α<br>(HNF1α-DNA focused)<br>(EMD-79882)<br>(PDB 39YB) |
|---|---|---|---|---|
|  | Class I<br>(EMD-79868)<br>(PDB 39XH) | Class II<br>(EMD-79878)<br>(PDB 39XX) |  |  |
| Magnification | 36kx | 36kx | 36kx | 36kx |
| Voltage (kV) | 200 | 200 | 200 | 200 |
| Electron exposure (e–/Å<sup>2</sup>) | 50 | 50 | 50 | 50 |
| Defocus range (µm) | -0.8 to -1.8 | -0.8 to -1.8 | -0.8 to -1.8 | -0.8 to -1.8 |
| Pixel size (Å) | 1.141 | 1.141 | 1.141 | 1.141 |
| Symmetry imposed | C1 | C1 | C1 | C1 |
| Initial particle images (no.) | 817,575 | 817,575 | 156,345 | 318,007 |
| Final particle images (no.) | 115,125 | 117,895 | 22,001 | 49,424 |
| Map resolution (Å)<br>FSC (0.143) | 3.5 Å | 3.7 Å | 6.2 Å | 4.3 Å |
| Map resolution range (Å) | 2.6-4.0 Å | 2.9-4.0 Å | 4.9-11.7 Å | 3.5-4.8 Å |
| Model resolution (Å)<br>FSC (0.5) | 3.8 | 4.2 | 7.1 | 5.0 |
| Map sharpening *B* factor (Å<sup>2</sup>) | -50 | -50 | -50 | -30 |
| Model composition<br>Non-hydrogen atoms<br>Protein residues<br>Nucleotides | 11943<br>733<br>298 | 11127<br>701<br>271 | 12016<br>786<br>282 | 3829<br>346<br>48 |
| *B* factors (Å<sup>2</sup>)<br>Protein<br>Nucleotides | 73.2<br>122.7 | 32.7<br>83.7 | 114.9<br>152.4 | 124.3<br>84.7 |
| R.m.s. deviations<br>Bond lengths (Å)<br>Bond angles (°) | 0.005<br>0.963 | 0.005<br>1.004 | 0.006<br>1.001 | 0.005<br>0.900 |
| Validation<br>MolProbity score<br>Clashscore<br>Rotamers outliers (%) | 1.4<br>6.3<br>0.33 | 1.5<br>8.3<br>0.34 | 1.96<br>16.5<br>0.77 | 1.7<br>16.7<br>0.7 |
| Ramachandran plot<br>Favored (%)<br>Allowed (%)<br>Outliers (%) | 98.2<br>1.8<br>0.0 | 98.3<br>1.7<br>0.0 | 96.4<br>3.4<br>0.2 | 97.9<br>1.6<br>0.5 |

